# lincRNA RP24-315D19.10 promotes endometrial decidualization via upregulation of hnRNPA2B1

**DOI:** 10.1101/2022.09.07.506883

**Authors:** Liping Tan, Rufei Gao, Xuemei Chen, Yanqing Geng, Xin Yin, Peng Chuan, Xinyi Mu, Yan Su, Yan Zhang, Fangfang Li, Junlin He

## Abstract

Sufficient decidualization is necessary to maintain successful pregnancy. The physiological function and underlying molecular mechanisms of intergenic long non-coding RNA (lincRNAs) in this process remain largely unknown. Herein, we identified a lincRNA, RP24-315D19.10, which is highly expressed during mouse decidualization during early pregnancy, by performing RNA-sequencing (RNA-seq) analysis and weighted gene co-expression network analysis (WGCNA). Detailed cell and molecular assays revealed that lincRNA RP24-315D19.10 knockdown blocked decidualization in primary mouse endometrial stromal cells (mESCs), suggesting that RP24-315D19.10 is a promoting factor for decidualization. Mechanistically, cytoplasmic RP24-315D19.10 directly interacted with heterogeneous nuclear ribonucleoprotein A2B1 (hnRNPA2B1) and upregulated its protein level. Moreover, we found that hnRNPA2B1 is involved in the regulation of decidualization through loss- and gain-of-function studies *in vitro*. Clinically, patients diagnosed with spontaneous miscarriage were found to have lower hnRNPA2B1 levels than healthy individuals, suggesting that RP24-315D19.10-regulated hnRNPA2B1 may participate in the development and progression of early spontaneous abortion. Our study indicates that RP24-315D19.10 enhances endometrial decidualization in a hnRNPA2B1-dependent manner, providing further insights into this physiological process.

## Introduction

Implantation of blastocysts into the maternal uterus is critical for mammalian reproduction (1). In humans, the rate of natural conception is limited to approximately 30% (2), and 75% of failed pregnancies are attributed to implantation failure (2, 3). Prior studies have demonstrated that the blastocyst’s state of activity is a critical determinant of implantation (4). However, emerging evidence has shown that appropriate priming of the endometrium, a process known as decidualization, may contribute more to implantation than the quality of the embryo (5, 6).

Decidualization refers to the transformation of the stromal compartment of the endometrium to accommodate pregnancy. Endometrial stromal cells proliferate extensively and differentiate into specialized secretory decidual cells. The initiation of uterine decidualization in mice is stimulated by blastocysts. However, in humans, initiation of this process is driven by a postovulatory rise in progesterone levels and increased local cAMP production (3, 7). Decidual cells can remodel the extracellular matrix essential for controlling embryo implantation into the uterine wall, providing nutrition, and acting as an immune barrier for subsequent placental establishment (5, 7, 8). In contrast, suboptimal or defective decidualization leads to aberrations in placentation and ultimately to adverse pregnancy outcomes (3, 6), such as severe preeclampsia (9–11), endometriosis-associated infertility (12). Previous evidence indicates that several signaling molecules (3, 13, 14), multiple transcription regulators (15–17), cell cycle regulators (18, 19) and epigenetic regulators (12) tightly govern the process of decidualization. However, to the best of our knowledge, the precise molecular mechanisms underlying embryo implantation, particularly decidualization, are still not fully elucidated.

Long intergenic noncoding RNAs (lincRNAs) are defined as RNAs longer than 200 nucleotides with an apparent lack of protein-coding potential. lincRNAs have been distinguished from a broader lncRNA class of transcripts, as the former do not physically overlap with any protein-coding genes (20, 21). It is well documented that lincRNAs, characterized by remarkable tissue specificity, have important biological functions (22–24). Their aberrant expression in diverse tissues is closely linked to diseases such as cancer, nervous system diseases, and metabolic diseases (25–27). lincRNAs can also modulate gene expression in cis and trans at the transcriptional and post-transcriptional levels(28–30). To date, only few lincRNAs have been reported to be involved in the process of decidualization (31); hence, the specific molecular mechanisms by which lincRNAs regulate decidualization remain poorly understood.

In this study, we first performed RNA-seq of the endometrium in mice at the implantation and inter-implantation sites during decidualization. Combined WGCNA, we yielded a lincRNA named RP24-315D19.10, which is associated with decidualization. We found that RP24-315D19.10 was significantly upregulated in the mouse endometrium at the implantation sites, artificially decidualized endometrial tissues, and mouse decidual stromal cells. We demonstrated the positive impact of RP24-315D19.10 on endometrial stromal cell decidualization. Mechanistically, we found that RP24-315D19.10 directly binds to hnRNPA2B1 and stimulates hnRNPA2B1 protein expression, thus facilitating endometrial decidualization. These results provide further insights into the role and regulatory mechanism of lincRNAs in decidualization.

## Results

### RNA-seq in mice endometrium at implantation and inter-implantation sites

We first profiled the lncRNAs and mRNAs from uterine tissues at the implantation and inter-implantation sites using RNA-seq (Figure 1A). As shown in Figure 1 (B and C), lincRNAs accounted for 61.3% of all lncRNAs without protein-coding potential. When criteria *P* < 0.05 and fold change ≥ 2.0 were accepted, 21 differentially expressed lncRNAs and 586 differentially expressed mRNAs were identified (Table 1, Figure 1, D and E, and Supplemental Figure 1, A and B). Combining Gene Ontology (GO) analysis and Kyoto Encyclopedia of Genes and Genomes (KEGG) analysis, we observed that the biological processes of cellular processes and cell cycle pathways were regulated by differentially expressed mRNAs (Supplemental Figure 1, C and D). Based on their effects on gene expression, lncRNAs are classified as follows: cis-acting lncRNAs (cis-lncRNAs) that regulate the expression of genes in close genomic proximity, and trans-acting lncRNAs (trans-lncRNAs) that regulate the expression of distant genes (29, 32). Thus, according to the positional relationship between lncRNAs and their target genes, the predicted target genes of all dysregulated lncRNAs can be broadly divided into two clusters: 1) co-location target genes, those within 100kb upstream and downstream, were regulated by cis-lncRNAs, and 2) co-expression target genes, highly correlated with the lncRNA expression profile (*r* > 0.95), were regulated by trans-lncRNAs. In GO analysis, co-location target genes were mainly clustered into negative regulation of biological processes and negative regulation of macromolecule metabolic process classification categories; co-expression target genes were mainly clustered into biological process and cellular process classification categories (Figure 1, F and H). KEGG analysis indicated that co-located target genes were involved in the alcoholism pathway and co-expression target genes were mainly involved in the cell cycle pathway (Figure 1, G and I). Furthermore, to verify the accuracy of the RNA-seq results, we randomly selected six dysregulated lncRNAs and assessed their relative expression levels in uterine tissues at implantation and inter-implantation sites. According to the RT-qPCR results, expressions of the six lncRNAs were consistent with those observed in RNA-seq (Supplemental Figure 2, A-F).

**Table.1.**
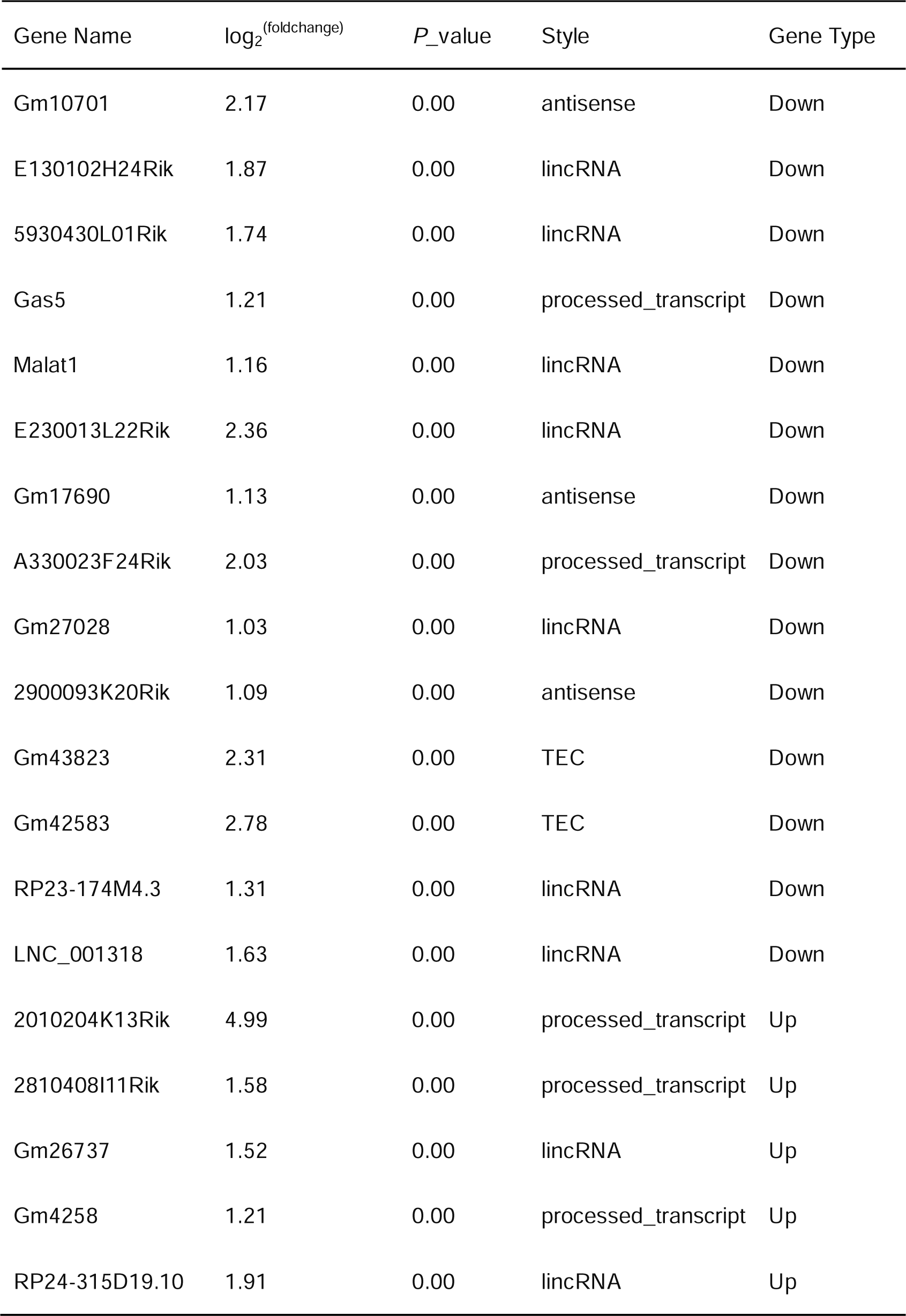
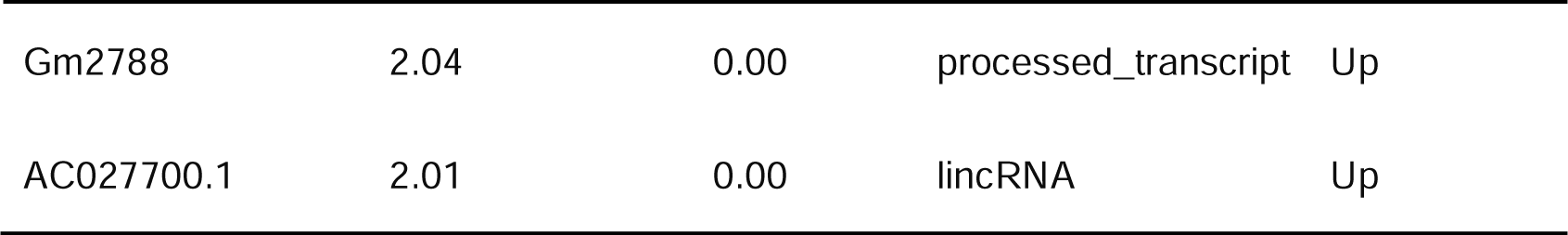
The 21 differentially expressed lncRNAs

**Figure 1.**
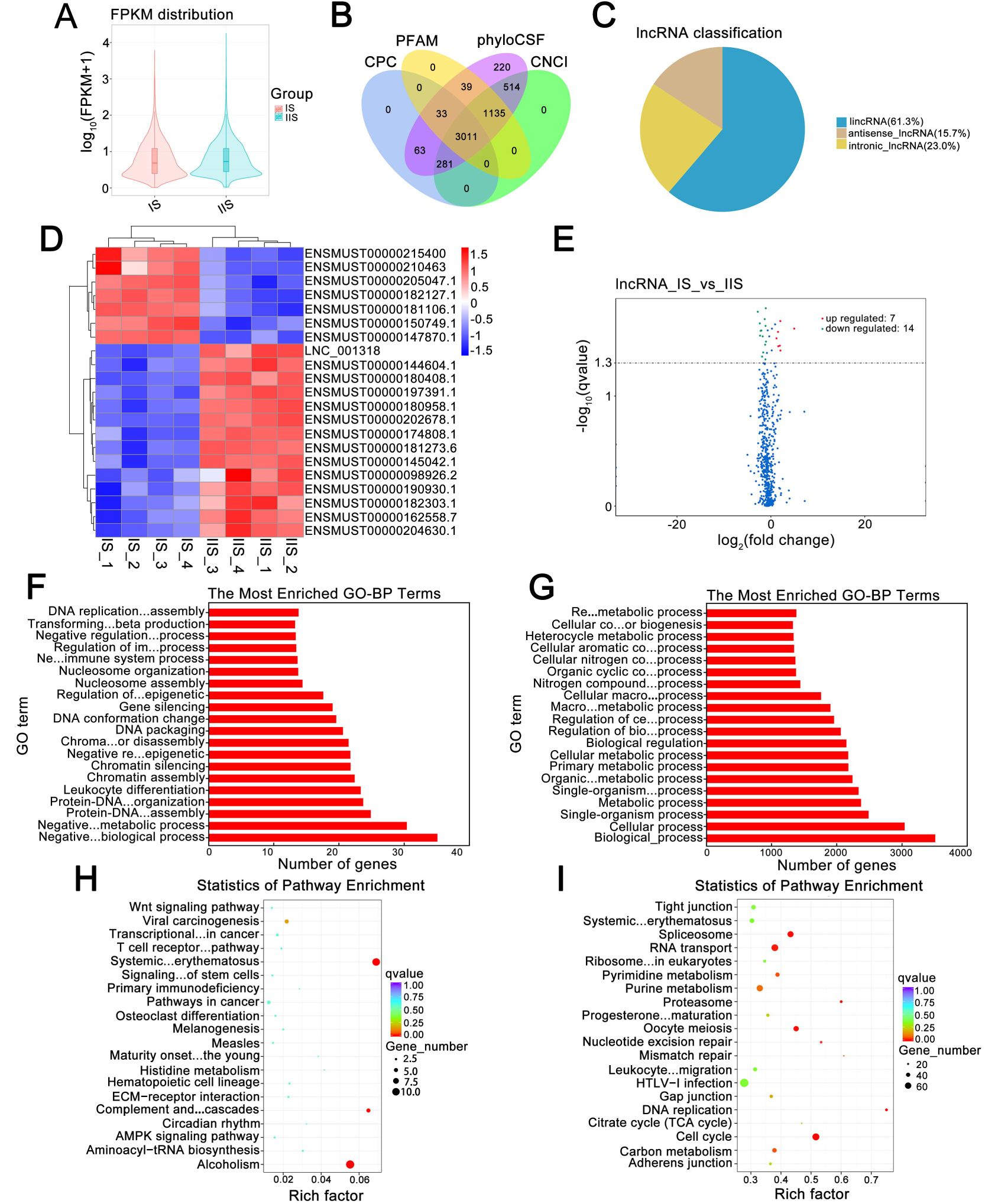
RNA-seq in mice endometrium at implantation sites and inter-implantation sites on the 6th day of pregnancy. (**A**): FPKM distribution between implantation sites and inter-implantation sites. The abscissa indicates sample name, and the ordinate indicates log ^(FPKM+1)^. (**B**): Venn diagram of identifying lncRNAs. Transcript coding potential screening was performed by CNCI, CPC, PFAM and phyloCSF, and finally 3011 lncRNAs were identified. (**C**): The pie chart shows classification of lncRNA, among them lincRNA accounts most (61.3%). (**D**): The cluster heat map shows lncRNAs with expression change fold ≥ 2 from RNA-seq data (*P* < 0.05). Red color indicates high expression level, and blue color indicates low expression level. (**E**): Volcano plot shows the distribution of differentially expressed lncRNAs. (**F**): Top 20 GO-BP terms enriched in co-location target genes. (**G**): Top 20 GO-BP terms enriched in co-expression target genes. (**H**): Top 20 KEGG pathways enriched in co-location target genes. (**I**): Top 20 KEGG pathways enriched in co-expression target genes. IS: implantation sites; IIS: inter-implantation sites.

### Identification of decidualization-associated hub lncRNAs using WGCNA

It is worth emphasizing that not all dysregulated lncRNAs play a role in decidualization. Therefore, there is an urgent need to establish relationships between lncRNAs and decidualization. Thus, we performed WGCNA, which provided systems-level insight to discover the lncRNA-decidualization relationship and to identify hub lncRNAs. Supplemental Figure 3A shows the cluster tree of all lncRNAs and mRNAs and the corresponding trait information. As shown in Supplemental Figure 3B, the soft-threshold power was set to five, and a topological matrix with non-scale features (scale-free R^2^ > 0.9) was obtained. Using the dynamic tree cut method, all transcripts were divided into 16 modules according to their correlations (Figure 2A). The heat map revealed the eigengene adjacency of the modules (Figure 2B).

**Figure 2.**
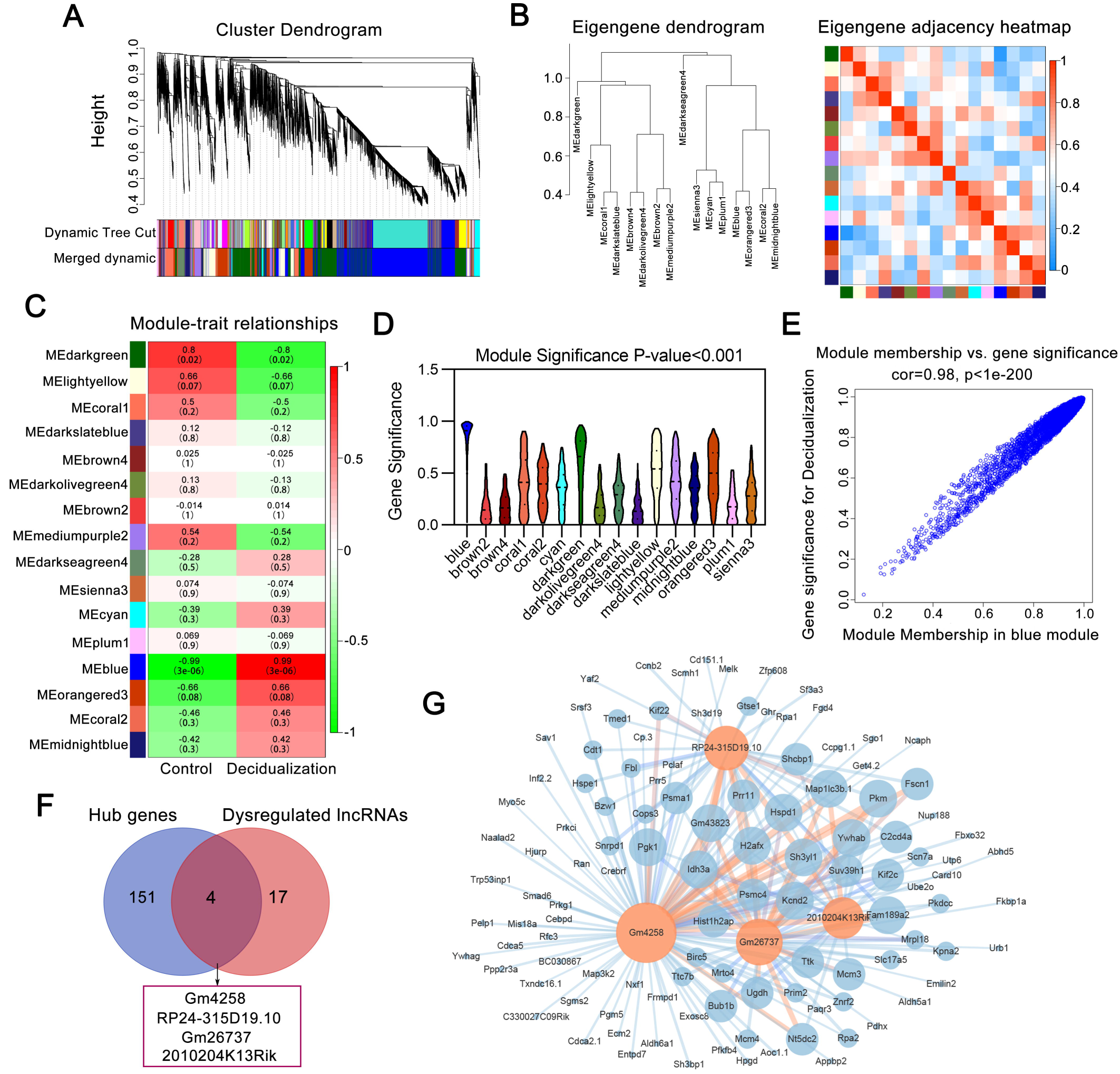
Identification of decidualization-associated hub lincRNAs by WGCNA. (**A**): Gene dendrogram obtained by hierarchical clustering of adjacency-based dissimilarity. The color row underneath the dendrogram shows the module assignment determined by the Dynamic Tree Cut. After merging the two modules with a correlation coefficient > 0.6, the final number of obtained modules was 16. (**B**): Hierarchical clustering dendrogram of modules eigengenes (MEs). Branches of the dendrogram group together eigengenes that are positively correlated. Each column and row in the heatmap correspond to one ME (labeled by color). In the heatmap, red indicates high adjacency (positive correlation), whereas blue indicates low adjacency (negative correlation). (**C**): The module–traitrelationships. Each column corresponds to a trait, and each row corresponds to a ME. The box is color-coded by correlation based on the color legend. The red indicates positive negative correlation and the green indicates negative correlation. The value in the box showed the correlation between ME and trait, the corresponding *P* values are shown below in brackets. (**D**): Bar plots of mean GS across modules. The higher the mean GS in a module is, the more significantly related the module is to decidualization. (**E**): Scatterplot of GS for decidualization (y-axis) vs. MM (x-axis) in blue module. (**F**): The Venn diagram displays the intersection between the 155 hub genes in the blue module and the 21 dysregulated lncRNAs identified by differential expression analysis. The 4 lncRNAs at the intersection of these two groups are listed in the Table below. (**G**): Top 200 co-expression connections of 4 lincRNAs and mRNAs in the blue module. In this network, the depth of color indicates the connectivity of a hub lincRNA. ME: module eigengenes; GS: gene significance; MM: module membership.

A module-trait relationship heatmap was created based on the Pearson correlation coefficients between module eigengenes (MEs) and trait information, and the blue module exhibited the strongest correlation with decidualization (*r* = 0.99, *P* < 0.001, Figure 2C). Moreover, the mean absolute gene significance (GS) values for decidualization were largest for the blue module (Figure 2D). In other words, the blue module exhibited the highest correlation with decidualization, and the correlation coefficient between GS and module membership (MM) was highest in the blue module (*r* = 0.98, *P* < 0.001, Figure 2E). Therefore, screening of hub RNAs was performed within the blue module, and 155 hub RNAs were identified as those with GS > 0.98 and MM > 0.98 (Supplemental Table 1). Hub lncRNA screening was required to meet these two conditions. First, the hub lncRNAs were part of the 155 hub RNAs. Second, the hub lncRNAs met the screening criteria for differentially expressed lncRNAs. Once a lncRNA met these two screening criteria simultaneously, it was defined as a hub lncRNA. As a result, a total of four hub lncRNAs, all of which can be categorized as lincRNAs, were identified (Figure 2F). Next, we constructed a lincRNA-mRNA co-expression network (top 200 according to weight) using Cytoscape 3.6.1 based on the mRNA expression profile in the blue module. As shown in Figure 2G, lincRNA Gm4258 had the highest connectivity, followed by RP24-315D19.10, suggesting their crucial role in decidualization.

### RP24-315D19.10 is upregulated during decidualization

The expression levels of the four hub lincRNAs, 2010204K13Rik, Gm26737, Gm4258, and RP24-315D19.10, in uterine tissues at implantation and inter-implantation sites on the 6th day of pregnancy were detected using RT-qPCR. In agreement with the RNA-seq results, the four hub lincRNAs were increased at the implantation sites compared to the inter-implantation sites (Figure 3A). To investigate the expression patterns of the four hub lincRNAs during decidualization, we established an artificially induced decidualization model. The significant increase in decidual/trophoblastic prolactin-related proteins (Dtprp) and unilateral uterine weight in the decidualized side versus the control side proved that our artificial decidualization model was successfully constructed *in vivo* (Figure 3, B-D). Using RT-qPCR, we observed significant upregulation of 2010204K13Rik and RP24-315D19.10, as well as downregulation of Gm26737 and Gm4258, in the decidualized side compared to the control side (Figure 3E). Furthermore, mESCs isolated from non-pregnant female mice were decidualized using 17β-estradiol and progesterone *in vitro*, and changes in their cytoskeletal organization were investigated using filamentous actin (F-actin) staining after artificially induced decidualization (Figure 3F). Compared to fibroblast-like mESCs, mouse decidual stromal cells (DSCs) had a typical polygonal shape with multiple nuclei and abundant actin stress fibers (Figure 3F). Additionally, the remarkable upregulation of Dtprp in DSCs proved that an *in vitro*-induced decidualization model was successfully constructed (Figure 3G). We next evaluated the above four hub lincRNAs in mESCs and DSCs and found that Gm26737, Gm4258, and RP24-315D19.10 exhibited increased expression in DSCs compared with mESCs. 2010204K13Rik was downregulated in the DSCs (Figure 3H). In summary, 2010204K13Rik, Gm26737, and Gm4258 were excluded owing to their inconsistent expression patterns. Thus, RP24-315D19.10 was selected as the candidate lincRNA for the follow-up study.

**Figure 3.**
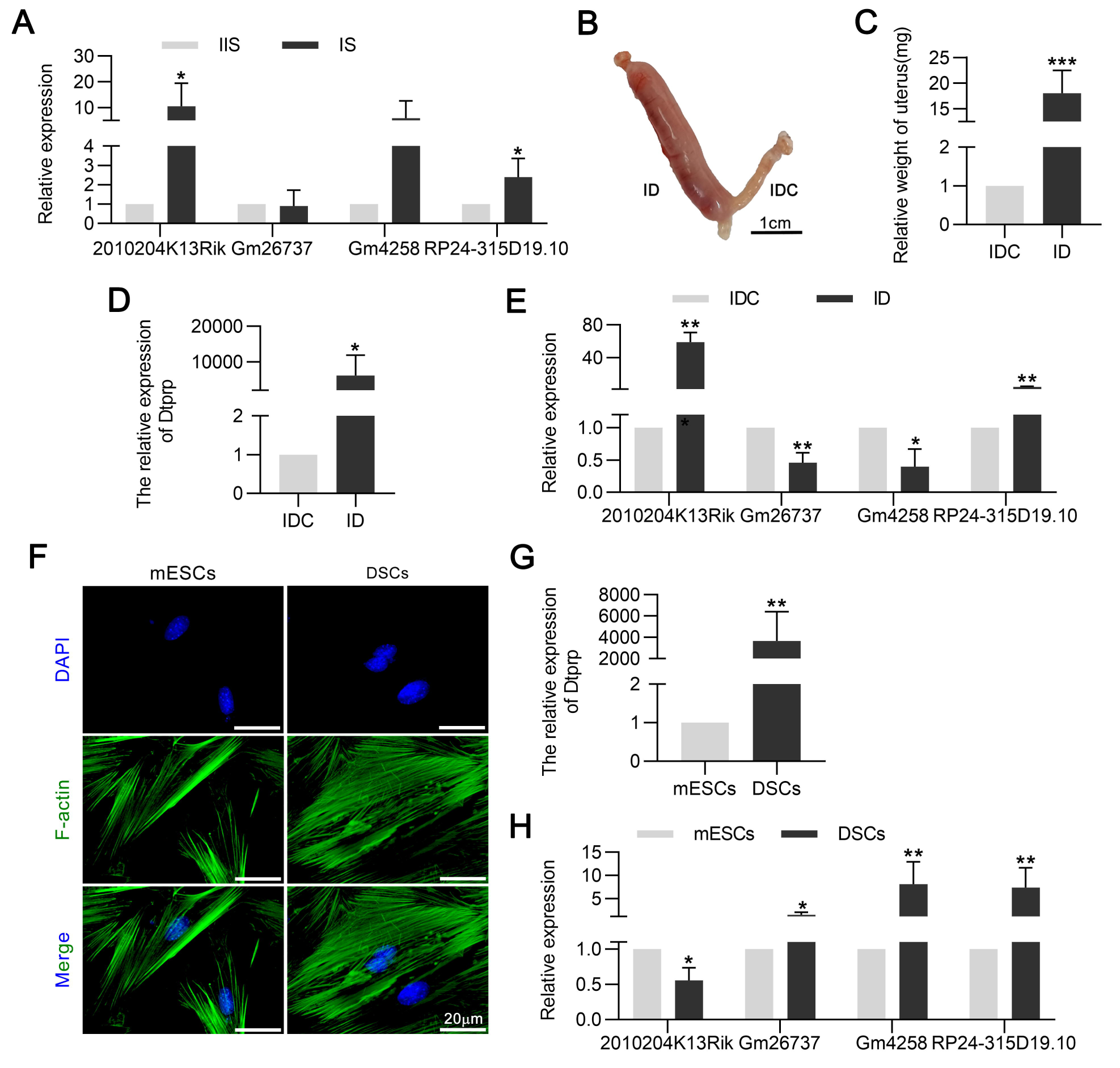
RP24-315D19.10 is upregulated during decidualization. (**A**): RT-qPCR detection of 4 hub lincRNAs expression at the IS and at the IIS on Day 6 of pregnancy (n=5). (**B**): A typical photograph showing the uterus after injecting oil on Day 8 of pseudopregnancy. (**C**): The comparison of bilateral uterine weight on Day 8 of pseudopregnancy (n=9). (**D**): Relative expression of Dtprp in endometrium derived from decidualized side and control side was measured by RT-qPCR (n=5). (**E**): Relative expression of above 4 hub lincRNAs in endometrium derived from decidualized side and control side was measured by RT-qPCR (n=5). (**F**): Fluorescent isothiocyanate labeled phalloidin was used to stain F-actin of DSCs and mESCs. (**G**): Relative expression of Dtprp in DSCs and mESCs was detected by RT-qPCR (n=4). (**H**): RT-qPCR measured the relative expression of 4 hub lincRNAs in DSCs and mESCs (n=4). \**P* < 0.05, \*\**P* < 0.01, \*\*\**P* < 0.001 compared with controls; data are presented as mean ± SD. IS: implantation sites; IIS: inter-implantation sites; Dtprp: decidual/trophoblast prolactin-related protein; IDC: control side, nothing injected; ID: oil side, artificially induced decidualization; mESCs: mouse endometrial stromal cells; DSCs: decidual stromal cells.

### RP24-315D19.10 promotes mESCs decidualization

RP24-315D19.10 (ENSMUST00000205047.1) is located on mouse chromosome 6: 48,860,329-48,866,083, and the transcript length of RP24-315D19.10 was 2194bp (Supplemental Figure 4A). The coding potential of RP24-315D19.10 was predicted using several prediction softwares, namely CPC, CNCI, PFAM, and phyloCSF, and the results showed that RP24-315D19.10 could not code any proteins (Supplemental Figure 4B). Owing to its specific and stable expression during decidualization, we investigated whether RP24-315D19.10 silencing resulted in the resistance of primary mESCs to decidualization *in vitro*. We transfected mESCs with three siRNAs targeting different sites within RP24-315D19.10 and the negative control, and RP24-315D19.10-silenced mESCs were subsequently decidualized using 17β-estradiol and progesterone for 24 h. Si-2 effectively knocked down RP24-315D19.10 expression, and was therefore used throughout this study (Figure 4A). Next, we examined the effect of RP24-315D19.10 silencing on morphological changes in mESCs during *in vitro* decidualization. As expected, si-2-treated mESCs failed to transform from a fibroblastoid-to-decidual phenotype (Figure 4B). Furthermore, RT-qPCR showed that RP24-315D19.10 depletion inhibited Dtprp expression (Figure 4C). The protein expression levels of Cox-2 and BMP2, marker molecules related to decidualization, were also significantly reduced after RP24-315D19.10 silencing (Figure 4D). The inseparable relationship between cell proliferation, apoptosis, and decidualization prompted us to explore whether RP24-315D19.10 knockdown affects mESCs proliferation or apoptosis. Thus, we measured the expression levels of proliferation-related factors Ki-67 and PCNA, as well as apoptosis-related factors Bax and Bcl2, by performing western blotting, immunofluorescence, and TUNEL staining. As indicated by the results, RP24-315D19.10 silencing significantly inhibited mESCs proliferation and promoted apoptosis during decidualization (Figure 4, E-G). These results suggest that RP24-315D19.10 promotes decidualization *in vitro*.

**Figure 4.**
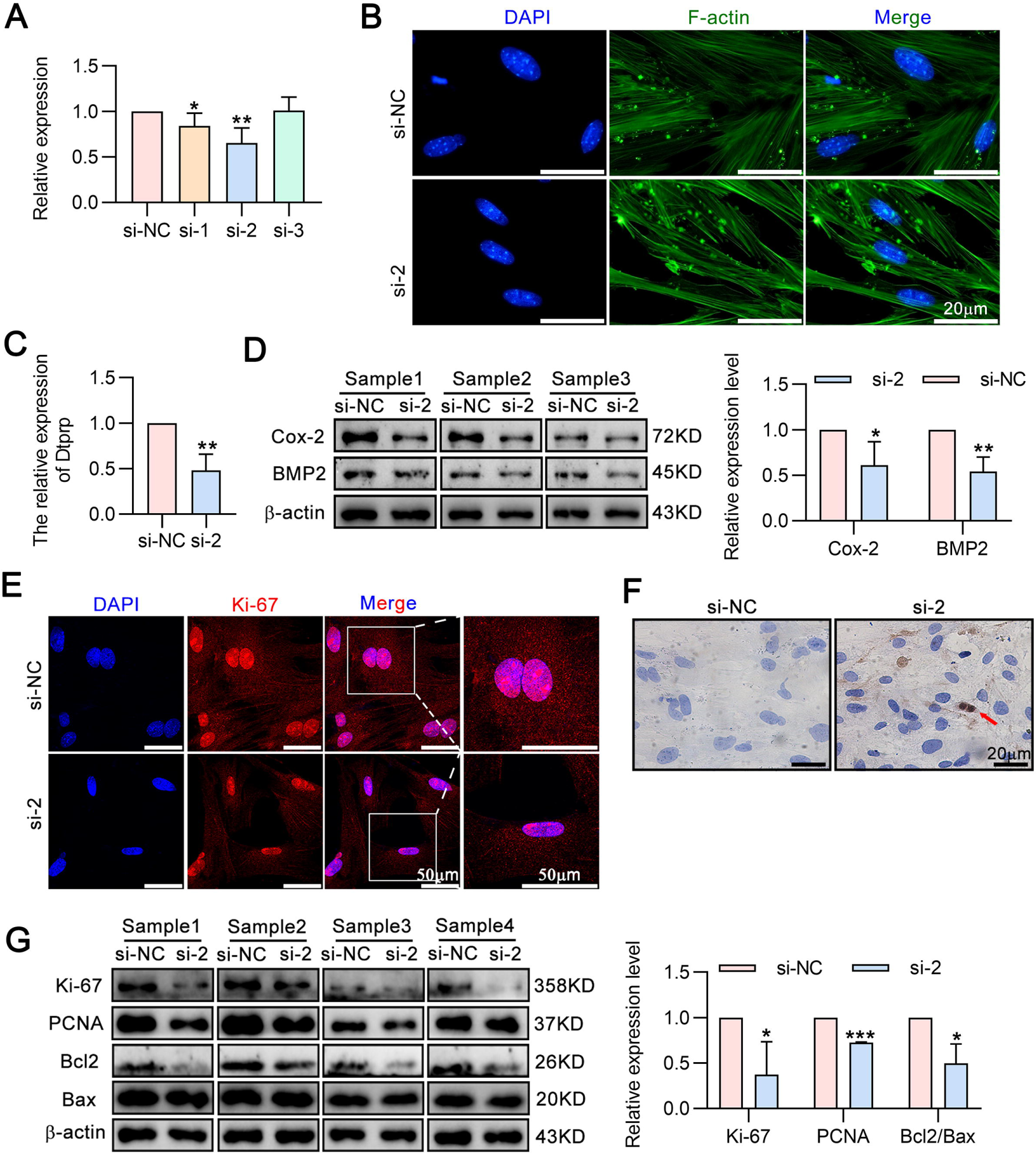
RP24-315D19.10 promotes decidualization *in vitro*. (**A**): The RP24-315D19.10 interference efficiency using 3 different siRNAs and negative control was detected by RT-qPCR (n=6). (**B**): Relative expression of Dtprp of DSCs transfected with si-2 or si-NC (n=4). (**C**): Fluorescent isothiocyanate labeled phalloidin was used to stain cellular F-actin of DSCs after RP24-315D19.10 silencing. (**D**): Western blot analysis indicates the expression of decidualization related markers after RP24-315D19.10 silencing (n=3). Histogram showing the quantification of the results for western blot. (**E**): Immunofluorescence of Ki-67 in DSCs transfected with si-2 or si-NC. (**F**): TUNEL method was used to observe the cell apoptosis after RP24-315D19.10 silencing. (**G**): Western blot analysis indicates the expression of proliferation related markers and apoptosis related markers after RP24-315D19.10 silencing (n=4). Histogram showing the quantification of the results for western blot. \**P* < 0.05, \*\**P* < 0.01, \*\*\**P* < 0.001 compared with controls; data are presented as mean ± SD. Dtprp: decidual/trophoblast prolactin- related protein; BMP2: bone morphogenic protein 2; Cox-2: Cyclooxygenase-2; PCNA: proliferating cell nuclear antigen; Bcl2: B-cell lymphoma 2.

### RP24-315D19.10 promotes decidualization via upregulation of hnRNPA2B1

Next, we sought to understand the underlying molecular mechanism by which RP24-315D19.10 regulates decidualization. We first used two online prediction tools: lncLocator (www.csbio.sjtu.edu.cn/bioinf/lncLocator/) and iLoc-LncRNA (lin-group. cn/server/iLoc-LncRNA/) to explore the intracellular localization of RP24-315D19.10, and we found that RP24-315D19.10 may be retained in the cytoplasm (Supplemental Figure 4, C and D). Next, we examined its subcellular localization using RNA fluorescence in situ hybridization (FISH) in both mESCs and DSCs (Figure 5A). The cytoplasmic localization of RP24-315D19.10 was also confirmed using a nuclear/cytoplasmic RNA separation assay. The assay revealed that RP24-315D19.10 was predominantly cytoplasmic (Figure 5B). In the cytoplasm, lncRNAs usually interact with diverse RNA-binding proteins, thereby regulating their expression(30). Therefore, to search for potential RP24-315D19.10–interacting proteins, we performed an RNA pull-down assay with a biotinylated RP24-315D19.10 probe, followed by mass spectrometry (Figure 5C). It was shown that 9 proteins based on the number of unique peptides ≥ 2 were found (Table 2). Moreover, based on our RNA-seq data, 1662 RP24-315D19.10 co-expression target genes were screened, with a > 0.95 correlation to its expression profile. As shown in the Venn diagram, RP24-315D19.10 co-expressed target genes shared three targets with nine proteins identified by RNA pull-down/MS (Figure 5D). Among them, hnRNPA2B1, a key member of the heterogeneous nuclear ribonucleoprotein family, was identified as a candidate because of its high abundance in immunoprecipitates pulled down by the biotinylated RP24-315D19.10 probe (Table 2).

**Table.2.**
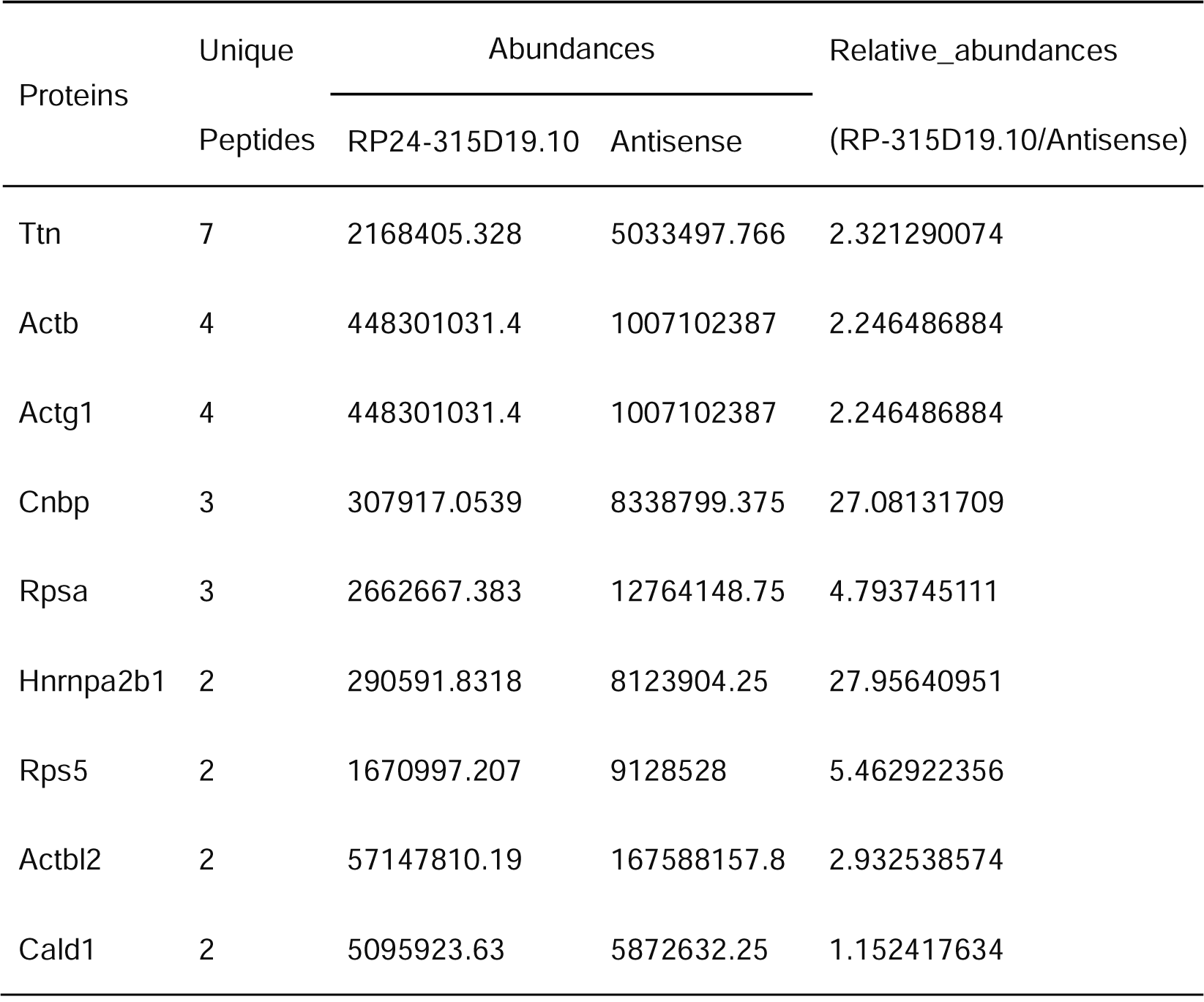
The 9 candidate interacting proteins of RP24-315D19.10 identified by mass spectrometry analysis

**Figure 5.**
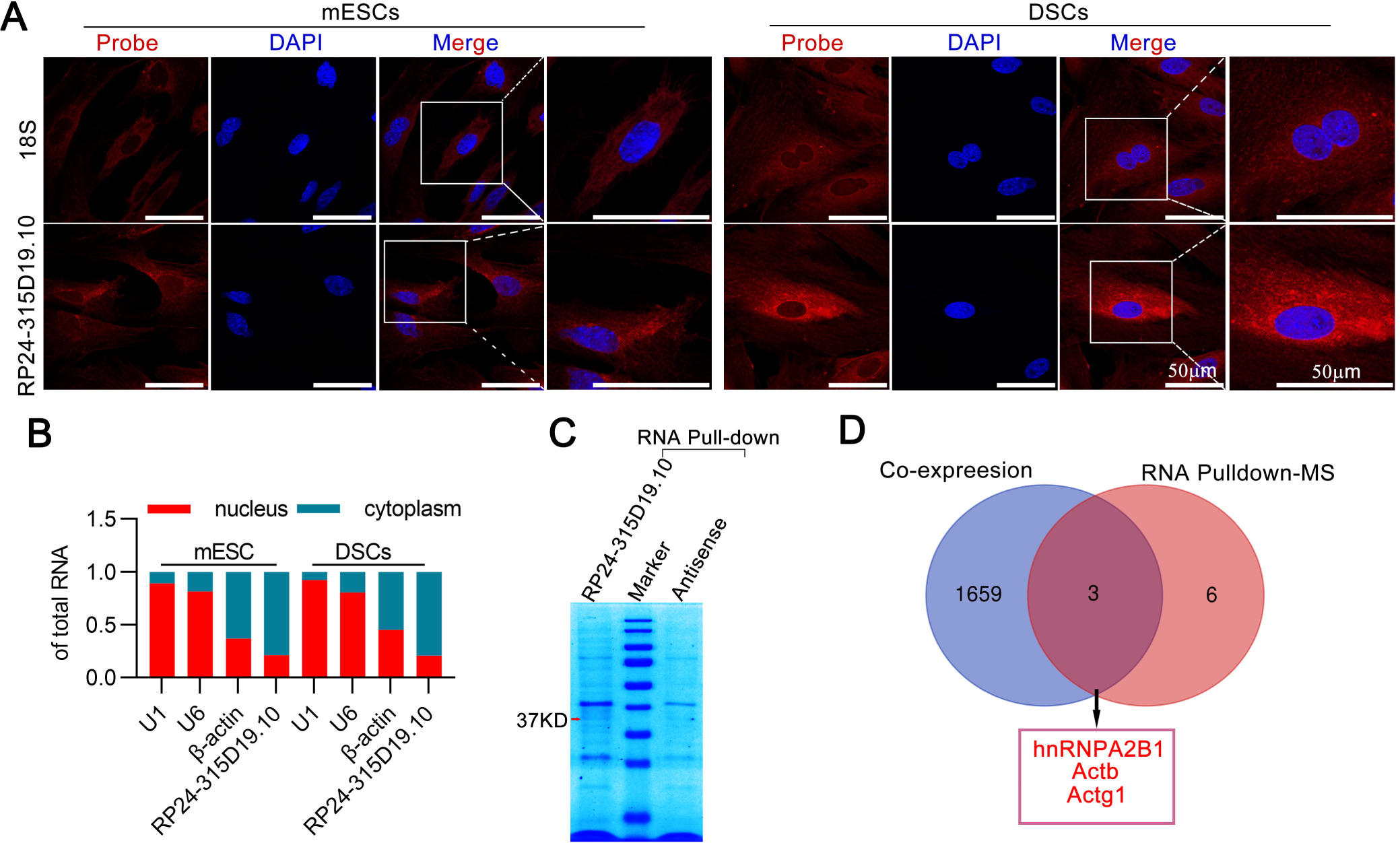
Screening the proteins interaction with cytoplasmic RP24-315D19.10. (**A**): Subcellular localization of RP24-315D19.10 and 18S in mESCs and DSCs detected by RNA FISH. (**B**): RP24-315D19.10 is enriched in cytoplasmic fraction of decidualized mESCs and DSCs. Expression levels of RP24-315D19.10, β-actin, U1 and U6 RNA in purified nuclear and cytoplasm fractions were measured by RT-qPCR. (**C**): Coomassie brilliant blue-stained SDS-PAGE gel of proteins immunoprecipitated from DSCs extracted by the biotin-labeled RP24-315D19.10 and the antisense RNA. The two lanes were used for mass spectrum determination and the arrows indicates hnRNPA2B1. (**D**): Venn diagram of identifying RP24-315D19.10 associated proteins. The 3 candidate proteins at the intersection of RP24-315D19.10-binding proteins detected by RNA pull-down/MS and co-expression target genes are listed in the bottom.

Next knocked RP24-315D19.10 down, and found that hnRNPA2B1 expression was significantly decreased in response (Figure 6A, Supplemental Figure 5A). At this point, we investigated whether there was a link between RP24-315D19.10-regulated hnRNPA2B1 and endometrial decidualization. We thus conducted hnRNPA2B1 knockdown experiments to confirm whether RP24-315D19.10 regulates mESCs decidualization *in vitro* and is also induced by hnRNPA2B1. Western blotting showed that si-A2B1-3 effectively knocked down hnRNPA2B1 expression in DSCs (Figure 6B and Supplemental Figure 5B). F-actin staining revealed that hnRNPA2B1 silencing had a significant negative effect on the morphological changes in mESCs during *in vitro* decidualization (Figure 6C). Consistently, hnRNPA2B1 silencing decreased Cox-2, BMP2, PCNA, and the Bcl2/Bax ratio (Figure 6D, Supplemental Figure 5C). Collectively, these results indicate a suppressive effect on decidualization in hnRNPA2B1 silencing DSCs. Given that both RP24-315D19.10 and hnRNPA2B1 regulate the process of decidualization of mESCs *in vitro*, we expressed hnRNPA2B1 in RP24-315D19.10 silencing mESCs to confirm whether RP24-315D19.10 facilitates mESCs decidualization by regulating hnRNPA2B1. Rescue experiments indicated that hnRNPA2B1 overexpression significantly rescued the inhibitory effect of RP24-315D19.10 silencing on mESCs decidualization (Figure 6, E and F, Supplemental Figure 5D). Collectively, our data suggest that RP24-315D19.10 promotes mESCs decidualization via hnRNPA2B1.

**Figure 6.**
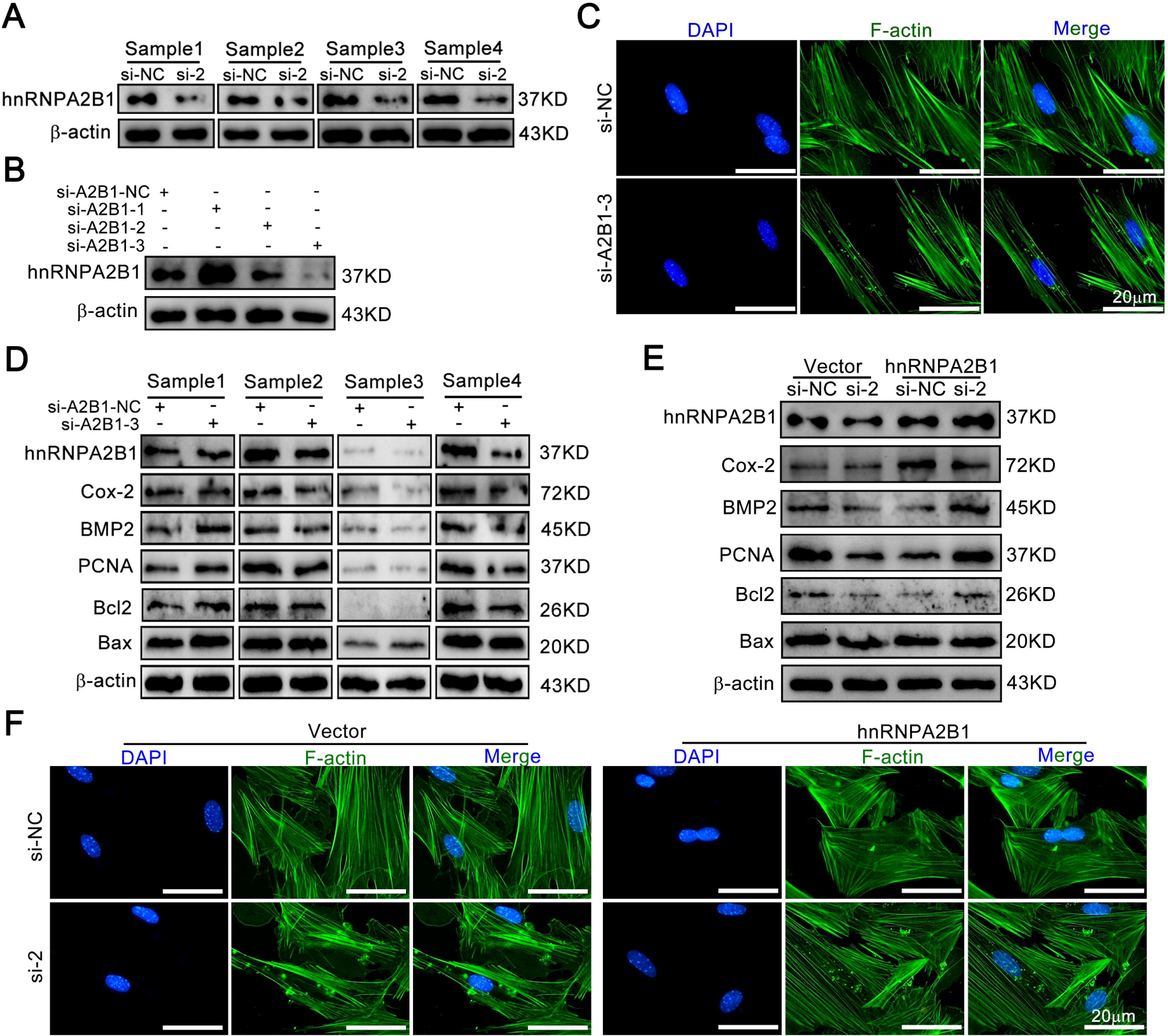
RP24-315D19.10 silencing inhibits hnRNPA2B1 promotion of decidualization *in vitro*. (**A**): Western blot analysis indicates the hnRNPA2B1 expression after RP24-315D19.10 silencing. (**B**): Western blot analysis indicates the hnRNPA2B1 expression after transfected with 3 different siRNAs targeting hnRNPA2B1 and negative control. (**C**): Fluorescent isothiocyanate labeled phalloidin was used to stain cellular F-actin of DSCs after hnRNPA2B1 silencing. (**D**): Western blot showed the abnormal expression levels of decidualization related markers, cell proliferation related markers and apoptosis related markers after knocking down hnRNPA2B1 *in vitro*. (**E**): After overexpressed hnRNPA2B1, the decidualization related markers, cell proliferation related markers and apoptosis related markers in RP24-315D19.10 silencing DSCs were detected by western blot. (**F**): After overexpressed hnRNPA2B1, the morphological changes in RP24-315D19.10 silencing DSCs were detected by F-actin staining. BMP2: bone morphogenic protein 2; Cox-2: Cyclooxygenase-2; PCNA: proliferating cell nuclear antigen; Bcl2: B-cell lymphoma 2.

However, more evidence is needed to confirm the interaction between RP24-315D19.10 and hnRNPA2B1. For this reason, proteins pulled down by the biotinylated RP24-315D19.10 probe were subjected to western blotting and further validated that RP24-315D19.10 could interact with hnRNPA2B1 directly (Figure 7A). Similarly, RIP-qPCR assay indicated that the hnRNPA2B1 antibody instead of the IgG antibody could successfully capture RP24-315D19.10 (Figure 7B). To further confirm the RP24-315D19.10/hnRNPA2B1 interaction, RNA-binding regions of hnRNPA2B1 protein and the binding sites of RP24-315D19.10 that interacted with hnRNPA2B1 protein were predicted using catRAPID [catRAPID(tartaglialab.com)], which contains catRAPID omics v2.0 module (33), catRAPID signature module (34) and others. The overall RNA-binding propensity of hnRNPA2B1 calculated using the catRAPID signature was 0.7 (Figure 7C). Next, catRAPID omics v2.0 module and catRAPID interactions with large RNAs module further revealed that there were two hnRNPA2B1-binding elements (hnRNPA2B1-BEs), the 11–258 and 1000–1249 nucleotide positions of the RP24-315D19.10 sequence, with a propensity of 0.79 (Figure 7D, Supplemental Figure 6, A and B). Furthermore, search results of the blast-like alignment tool (BLAT) of the UCSC Genome Browser(35) revealed that human lincRNA TRAM2-AS1, linc00276, and linc01517 share homologous sequence with RP24-315D19.10 (Figure 7E). Among the 16523 predicted hnRNPA2B1 binding human lncRNAs, TRAM2-AS1, linc00276 and linc01517 were predicted by the catRAPID omics v2.0 module to interact with hnRNPA2B1(Figure 7F). Based on binding site prediction and homology comparison analysis, we presumed that the hnRNPA2B1 protein interacted selectively with the ∼-142ccccc∼-167 region of the RP24-315D19.10 sequence. To determine whether the binding of hnRNPA2B1 was dependent on the region on RP24-315D19.10, the ∼-142ccccc∼-167 region was mutated (Figure 7G). We measured the dissociation constants (*KD*) of RP24-315D19.10-hnRNPA2B1 using biolayer interferometry (BLI) analysis and found that mutation of the ∼-142ccccc∼-167 region decreased the *KD* of RP24-315D19.10-hnRNPA2B1 (Figure 7H), suggesting that the RP24-315D19.10-hnRNPA2B1 interaction was significantly reduced. These results indicate that the ∼-142ccccc∼-167 region on RP24-315D19.10 is critical for the binding of hnRNPA2B1.

**Figure 7.**
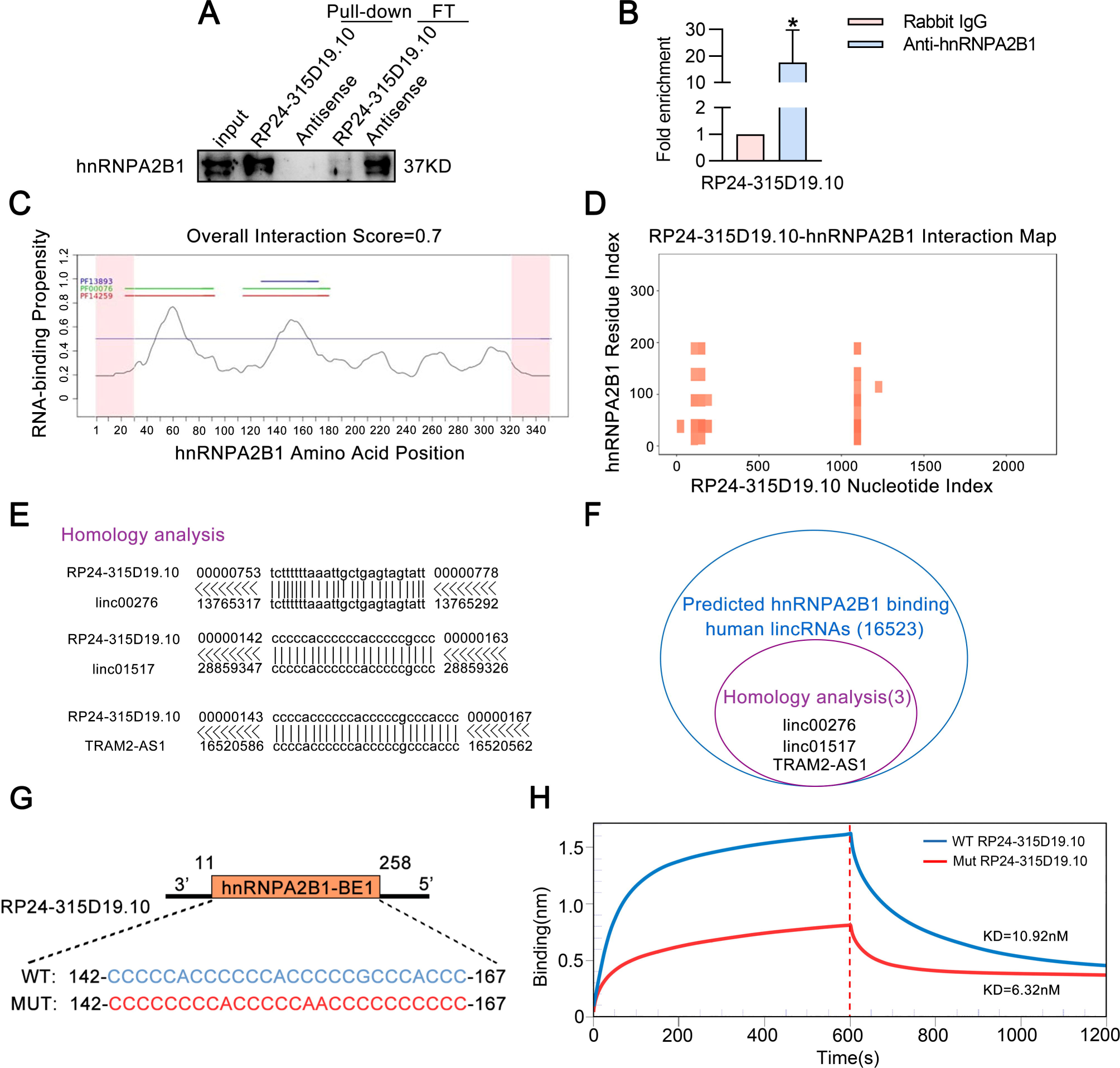
Recognition of RP24-315D19.10-hnRNPA2B1 binding regions. (**A**): Western blot showed that hnRNPA2B1 was enriched in the protein complexes pulled down by RP24-315D19.10. Pull-down indicates the RNA binding proteins. FT represents the flow-through proteins after RNA–protein binding reaction. Input indicates the total cell proteins used. (**B**): RIP assay detection of the interaction between hnRNPA2B1 and RP24-315D19.10 by using a hnRNPA2B1 antibody or isotype-matched control IgG, and the amount of RP24-315D19.10 in the precipitates was analyzed by RT-qPCR. (**C**). CatRAPID signature module prediction of the RNA-binding propensity for hnRNPA2B1 protein followed by prediction of RNA-binding regions. Overall interaction scores above 50% indicate propensity to bind. (**D**): CatRAPID omics v2.0 module prediction of the interaction map between hnRNPA2B1 protein and RP24-315D19.10. (**E**): UCSC database showed that RP24-315D19.10 has homologous sequence with 3 human lincRNAs. (**F**): The CatRAPID omics v2.0 module predicts that hnRNPA2B1 binds to 16523 human lncRNAs, including the 3 lincRNAs with homologous sequences to RP24-315D19.10. (**G**): Schematic of substitutions mutations of the ∼-142ccccc∼-167 region on RP24-315D19.10 sequence. (**H**): The interaction between *in vitro* transcribed RP24-315D19.10 and hnRNPA2B1 protein was detected by BLI binding affinity assay. WT: wild type RP24-315D19.10; MT: mutant RP24-315D19.10. \**P* < 0.05 compared with controls; data are presented as mean ± SD.

### hnRNPA2B1 expression in human endometrium

According to homology analysis using the BLAT program, the sequence of mouse hnRNPA2B1 has 95.7% homology with that of humans. Furthermore, hnRNPA2B1 was highly enriched in healthy human uteri (Supplemental Figure 7A), as determined using the online database GTEx (Genotype-Tissue Expression, http://www.gtexportal.org/home/). In humans, the uterus is considered receptive during the mid-secretory phase, in which human endometrial stromal cells are transferred into decidual cells (decidualization) (1). To validate whether hnRNPA2B1 is associated with human decidualization, we examined its expression during the normal human menstrual cycle. In the GEO database (accession no. GSE6364 at www.ncbi.nlm.nih.gov/geo) (36), a significant increase in hnRNPA2B1 mRNA levels in the human endometrium was observed gradually with the menstrual cycles (Figure 8A). Furthermore, hnRNPA2B1 was positively related to the prolactin receptor (PRLR) (Figure 8, B-D), which is considered a human decidualization marker. hnRNPA2B1 protein levels in decidual tissues derived from healthy volunteers and patients diagnosed with spontaneous abortion were analyzed using western blotting and immunohistochemistry.

**Figure 8.**
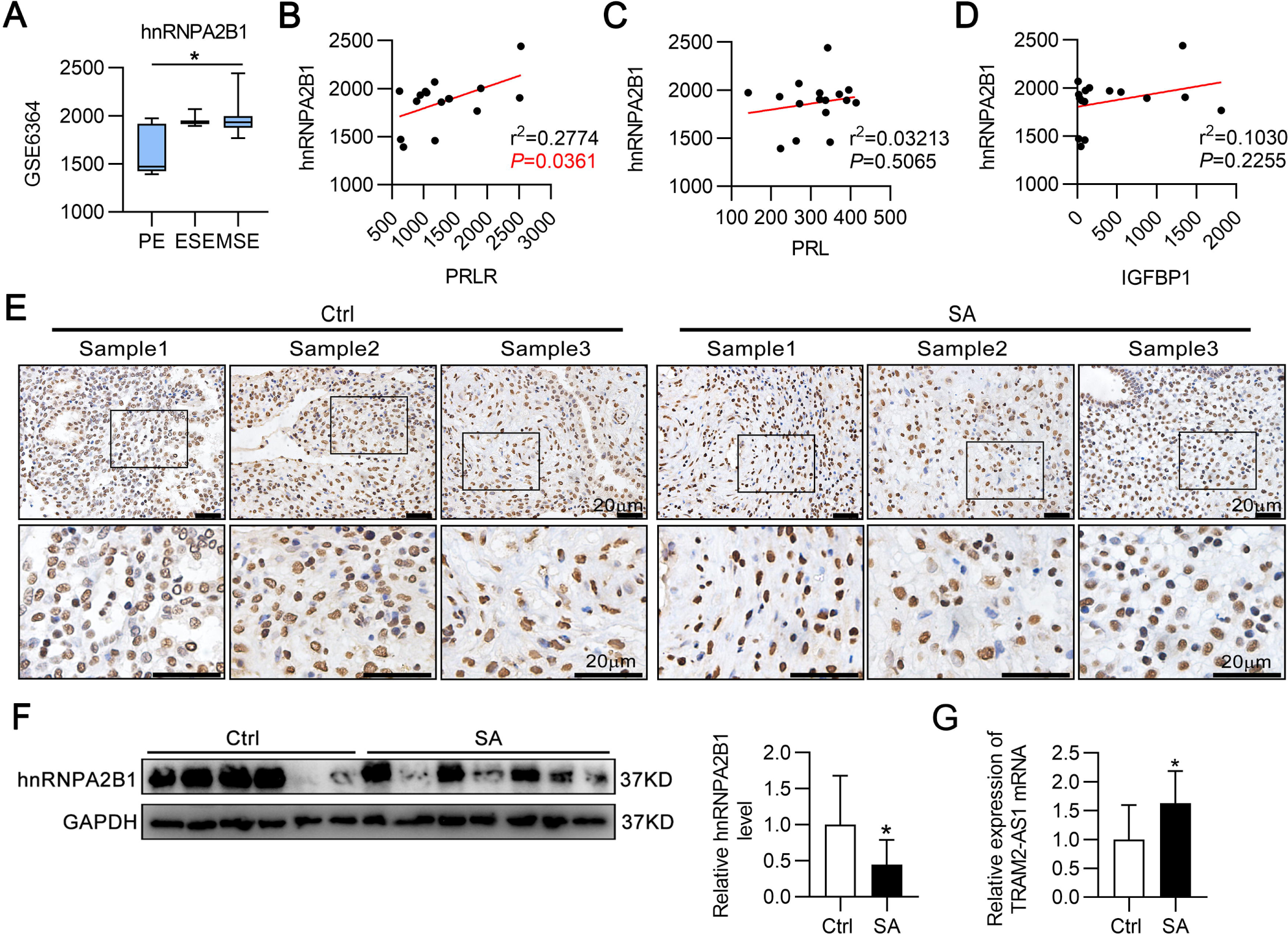
hnRNPA2B1 expression in human endometrium. (**A**): hnRNPA2B1 expression profile in human normal endometrium at distinct phases of the menstrual cycle. Corresponding data were obtained from GEO database (GSE6364). (**B**): Scatter plots displaying the correlation between hnRNPA2B1 mRNA expression and levels of PRLR, (**C**) PRL, and (**D**) IGFBP1. (**E**): Immunohistochemistry staining of hnRNPA2B1 in the decidual tissues of women (n=3). (**F**): Western blot showed the hnRNPA2B1 protein expression level in the Ctrl group (n=6) and SA group (n=7). Histogram showing the quantification of the results. (**G**): RT-qPCR showed lincRNA TRAM2-AS1 expression level in the Ctrl group (n=6) and SA group (n=7). Ctrl: Healthy volunteers (legal termination). SA: Patients diagnosed with spontaneous abortion. PE: Proliferative; ESE: Early Secretory; MSE: Mid-secretory. PRLR: Prolactin receptor. \**P* < 0.05 compared with controls; data are presented as mean ± SD.

The results showed that hnRNPA2B1 protein levels in the spontaneous abortion group were significantly lower than those in the healthy control group (Figure 8, E and F). Moreover, the level of the human lincRNA TRAM2-AS1, which showed some sequence conservation with RRP24-315D19.10, was significantly higher in the spontaneous abortion group than in the normal control group (Figure 8G), suggesting that TRAM2-AS1 is associated with spontaneous abortion. These results indicated that RP24-315D19.10-regulated hnRNPA2B1 may be involved in human decidualization.

## Discussion

It is widely known that only a small minority of lncRNAs’ functions and mechanisms of action have been well studied, despite thousands of lncRNA genes being identified. Among these deeply studied lncRNAs, many function in specific interactions with other cellular factors, such as proteins, DNA, and other RNA molecules (28). Several studies have shown that lncRNAs participate in the regulation of human disease pathogenesis through diverse mechanisms and serve as potential diagnostic and therapeutic targets (20, 37). To date, several lncRNAs associated with endometrial decidualization have been identified, including TUNAR(38), HOXA11-AS(39), Hand2os1(40), and PGK1P2(41). Studies have shown that these lncRNAs are abnormally expressed during endometrial decidualization in early pregnancy, implying that lncRNAs may play an important role in the development of endometrial decidualization.

In our study, we sought to identify lncRNAs associated with endometrial decidualization using RNA-seq analysis of uterine tissues at the implantation and inter-implantation sites of mice during early pregnancy. The present study found that the dysregulated lincRNA RP24-315D19.10 identified using WGCNA, which could not only find hub genes from complex gene expression data but also establish the relationships between hub genes and clinical traits (42, 43), might play a crucial role in decidualization. Next, we highlighted the function and mechanisms of RP24-315D19.10 in regulating endometrial decidualization. We demonstrated that RP24-315D19.10, which is significantly upregulated during decidualization, can enhance mESCs decidualization *in vitro* using loss-of-function experiments. This can be evidenced by reduced morphological changes and downregulation of Ki-67, PCNA, Bcl2/Bax, Cox-2, and BMP2 in DSCs after RP24-315D19.10 silencing. Together with the published report of human endometrial stromal cells decidualization-related lincRNA LINC00473(31), our study confirms the regulatory roles of lincRNAs in decidualization. Knockdown of RP24-315D19.10 resulted in mESCs failing to respond to decidualization induction *in vitro*, highlighting the positive effect of RP24-315D19.10 on decidualization.

We further showed that RP24-315D19.10 may serve as a trans-acting factor to modulate hnRNPA2B1 protein levels in a post-transcriptional manner. hnRNPA2B1, a key member of the heterogeneous nuclear ribonucleoprotein family, is an RNA-binding protein that localizes in both the cytoplasm and nucleus (44). hnRNPA2B1 can bind to a specific domain of RNA, thereby coordinating RNA levels through multiple mechanisms including transcriptional regulation, alternative splicing, transport, promotion of mRNA stabilization or degradation, and translation regulation(45). However, previous studies have shown that hnRNPA2B1 is also regulated by lncRNAs. For example, linc01232 directly interacts with hnRNPA2B1, which stabilizes hnRNPA2B1 by inhibiting ubiquitin-dependent degradation, subsequently facilitating pancreatic cancer progression (46). Another notable example is the lncRNA miR503HG, which is a hepatocellular carcinoma that negatively regulates lncRNAs. An investigation suggested that miR503HG could specifically interact with hnRNPA2B1 and promote its degradation via the ubiquitin-proteasome pathway (47). In this study, we observed a physical interaction between cytoplasmic RP24-315D19.10 and hnRNPA2B1 and decreased hnRNPA2B1 protein expression after RP24-315D19.10 silencing. It remains unknown whether hnRNPA2B1 mediates RP24-315D19.10 expression, thus warranting further investigation. The role of hnRNPA2B1 in cancer has received considerable attention in recent years, and hnRNPA2B1 may be used as a prognostic biomarker and therapeutic target (48, 49). However, in the field of implantation, particularly in endometrial decidualization, few studies have focused on hnRNPA2B1 expression. To the best of our knowledge, this is the first study to reveal that hnRNPA2B1 may play an essential role in endometrial decidualization. Similar to RP24-315D19.10 silencing, hnRNPA2B1 knockdown also led to decidualization defects in mESCs. Additionally, hnRNPA2B1 overexpression in mESCs offset the negative effect of RP24-315D19.10 silencing on mESCs proliferation or apoptosis and offset aberrant decidualization induced by RP24-315D19.10 silencing. These data indicate that RP24-315D19.10 enhanced mESCs decidualization through the hnRNPA2B1-dependent pathway.

Previous studies have found that impaired decidualization may increase the probability of miscarriage (50–53). Searching for hnRNPA2B1 function in human endometrial decidualization, we further revealed a significant increase in hnRNPA2B1 mRNA levels in the middle of the human menstrual cycle as well as a decrease in hnRNPA2B1 protein levels in spontaneous abortion, suggesting that hnRNPA2B1 may play a role in human decidualization. However, our research was limited to uncovering hnRNPA2B1 expression levels in human decidual tissues during spontaneous abortion. Therefore, additional functional investigations of hnRNPA2B1 in human decidualization are necessary.

Moreover, previous studies have implicated that lncRNAs can interact with proteins by binding to RNA sequence motifs to form specific lncRNA-protein complexes (lncRNPs), which participate in the post-transcriptional regulation of genes or signaling pathways(30). The 1093–1174 and 1249–1369 nucleotide positions of the PCAT1 sequence may bind to the FKBP51 protein, perturb the PHLPP/FKBP51/ IKKα complex, and activate AKT and NF-κB signaling, thus promoting the progression of prostate cancer(54). In addition to binding to certain sequence motifs, lncRNAs can fold into specific structures that can interact with proteins. For example, the lncRNA NKILA can interact with NF-B/I κB through two hairpin structures within nt 300–500, named hairpin A (nt 322–359) and hairpin B (nt 395–418), respectively, to form a stable NKILA:NF-κB:IκBα complex, thereby suppressing breast cancer metastasis (55). In this study, the potential functional regions of the RP24-315D19.10 sequence were found to interact with the hnRNPA2B1 protein using the catRAPID database and Biolayer interferometry analysis.

Although lincRNAs exhibit less cross-species conservation than protein-coding genes and other non-coding RNAs, they are not evolutionarily neutral and may have transcript-dependent and species-specific functions (20, 28). For instance, Ulitsky et al. identified a conserved 67 nt site of the lincRNA Cyrano, which is required for proper embryonic development in zebrafish, matched with human and mouse lincRNAs (56). Another lincRNA Gas5 contains conserved sequence-specific SR protein interaction domains that ensure a high level of the SR–Gas5 lincRNA complex, which regulates steroid-mediated transcriptional regulation, growth arrest, and apoptosis (57). Like Gas5, RP24-315D19.10 binds to hnRNPA2B1 protein through its potentially functional regions, which are homologous to the human lincRNA TRAM2-AS1. Intriguingly, the catRAPID database prediction results showed that TRAM2-AS1 may also bind to hnRNPA2B1 protein in a sequence-specific manner. Although we have not identified whether TRAM2-AS1 plays a role in human endometrial decidualization or the exact regulatory mechanism, we cannot entirely rule out the existence of the TRAM2-AS1-hnRNPA2B1 complex in decidualization regulation.

In conclusion, our findings suggest that mESCs decidualization is mediated by lncRNA RP24-315D19.10, which promotes hnRNPA2B1 expression. Nevertheless, further investigation is needed to establish a conclusive association between the RP24-315D19.10-hnRNPA2B1 complex and endometrial decidualization, as well as to ensure a deeper understanding of the molecular mechanisms involved.

## Materials and Methods

### Animals and treatments

Female and male CD1 mice of 8-week-old were purchased from Beijing Vital River Laboratory Animal Technology Company and maintained in Animal Facility of Chongqing Medical University. Female mice were mated with male mice to induce pregnancy and the females with copulatory plug in the next morning were considered as Day 1 of pregnancy (D1). Mice were sacrificed on D6, and all the endometrium at implantation and inter-implantation sites were collected for follow-up experiments. To establish the artificial induced decidualization model *in vivo*, Kunming mice, purchased from the Animal Center of Chongqing Medical University, were treated referred to previous study(58). Briefly, estrus female mice were mated with vasectomized male mice to induce pseudopregnancy and the females with copulatory plug in the next morning were considered as Day 1 of pseudopregnancy (PD1). At 9:00 am on PD4, unilateral uterus was injected with corn oil to induce decidualization, while the contralateral uninjected uterus served as a control. Pseudopregnant female mice were sacrificed on PD8. All the uteri were collected, part of them were stored at -80°C for western blot and RT-qPCR analysis, and the rest were fixed in 4% paraformaldehyde.

### RNA-seq analysis

Total RNA at implantation and inter-implantation sites was extracted using TRIzol reagent (Thermo Fisher Scientific). RNA-seq was done by Novogene Company (China) using standard Illumina protocols. Differentially expressed mRNAs and lncRNAs were screened as fold change ≥ 2.0 as well as *P* < 0.05, which was computed as the FPKM ratios of implantation sites over inter-implantation sites.

*Weighted gene co-expression network analysis.* To find out the key decidualization-related modules and hub lncRNAs, we constructed a weighted co-expression network based on our RNA-seq data using the R package “WGCNA”(42). We set soft-threshold power as five, which makes the constructed network conform to the power-law distribution. Dynamic Tree Cut method was employed to categorize lncRNAs and mRNAs with similar expression patterns into different modules. Each module was summarized by module eigengene (MEs) and was colored differently for visualization. The module with the highest correlation with decidualization was selected to screen hub lncRNAs. Hub RNAs were identified as those with gene significance (GS) > 0.98 and module membership > 0.98(MM). Cytoscape 3.6.1 was used to present the lncRNA-mRNA network (https://cytoscape.org/)(59).

### Isolation of primary mouse endometrial stromal cells and treatments

The primary mESCs were isolated as previously described(60). Mouse uterus tissues were minced, digested in trypsin (Sigma) for 2 h at 4 °C, followed by 30 min at 37 °C. After centrifugation and washing, tissues were digested in collagenase (Gibco) for 30 min at 37 °C. The digested tissues were passed through 70 μm strainer to obtain mESCs. Cells were then cultured in Dulbecco’s modified Eagle’s medium F-12 (Sigma) supplemented with 20% fetal bovine serum (Biological Industries), penicillin, streptomycin and amphotericin. After culture for 1 h, the medium was changed to remove unattached cells. The next day, the mESCs were induced for *in vitro* decidualization with 10 nmol/L 17 β-estradiol (Sigma) and 1 nmol/L Progesterone (Sigma) for 72 h.

### Total RNA extraction and Real-time quantitative PCR

The total RNA was isolated by TRI Reagent (Sigma). Reverse transcription was performed by using PrimerScript^TM^ RT Master Mix Kit (TaKaRa) according to directions. TB Green Premix Ex Taq^TM^ Ⅱ Kit (TaKaRa) was applied in determining the abundance of mRNA according to the manufacturer’s directions. The relative expression of mRNA was calculated by making use of 2^-ΔΔCt^ method and all results were normalized to β-actin. Primers sequences were detailed in Supplemental Table 3.

### Plasmid, siRNA and transfection

siRNA oligonucleotides with specificity for lincRNA RP24-315D19.10 and hnRNPA2B1 were designed and synthesized by GenePharma company (China). The hnRNPA2B1 plasmid was purchased from GenePharma company. All constructs were transiently transfected into mESCs using Lipofectamine 2000 (Invitrogen) according to the manufacturer’s protocol. At 6 hours post-transfection, mESCs were induced for *in vitro* decidualization for 24 h before further treatment. The siRNA sequences used in this study are listed in Supplemental Table 4.

### F-actin staining in vitro assay

mESCs and DSCs were washed with PBS for three times and fixed with 4% fresh para-formaldehyde for 15 min at room temperature, followed by incubation with FITC-phalloidin (Sigma) for 40 min at 37 °C in the dark. After repeated washing for three times, cells were stained with DAPI for 10 min at room temperature. Fluorescence microscope was employed to gain micrographs of cells.

### Western blot

Uterine tissues and cells were lysed in RIPA buffer (Beyotime Biotechnology). Proteins were separated by 10% SDS-PAGE gel and transferred onto 0.45 μm PVDF membranes. After blocking with 5% milk for 80 min at 37 °C, membranes were incubated overnight at 4 °C with different primary antibodies: rabbit monoclonal anti-BMP2 antibody (1:500; Abcam), rabbit monoclonal anti-Cox2 antibody (1:1000; Cell Signaling Technology), mouse monoclonal anti-PCNA (1:2000; Cell Signaling Technology), rabbit monoclonal anti-Ki-67(1:300; Bioss), rabbit monoclonal anti-Bax (1:1000; Cell Signaling Technology), rabbit monoclonal anti-Bcl2 (1:1,000; Cell Signaling Technology), rabbit polyclonal anti-hnRNPA2B1 (1:5000; Abcam) and mouse monoclonal anti-β-Actin (1:1000; Zhongshan Biosciences Inc.). The next day, membranes were incubated with HRP-conjugated secondary antibodies for 1h at 37°C. The protein bands were detected using chemiluminescence with imaging system (Bio-Rad). Quantity One version 4.6.2 (Bio-Rad) was used to quantify each band and then convert the graphical results into digital results. β-Actin was used as an internal reference.

### Immunocytochemistry

Transfected cells were fixed in 4% paraformaldehyde for 15 min, permeabilized with 0.5% Triton X-100 for 15 min and blocked with goat serum for 15 min. Cells then were incubated at 4 °C overnight with Anti-Ki-67 antibody (1:500; Bioss), followed by further incubation at 37 °C for 1 h with rabbit IgG. DAPI was used to visualize nuclei. Confocal microscopy was used to observe the image.

### Isolation of nuclear and cytoplasmic RNA

Nuclear and cytoplasmic RNA separation assay was conducted using RNA Subcellular Isolation Kit (Active motif), according to previously detailed method(61). Next, the RNA extracted from nuclear and cytoplasmic fractions was for the following RT-qPCR analysis. Results were analyzed by 2^−ΔCt^ [2^−(NUC Ct−CYT Ct)^] method (ΔCt method) as previously described(62). The primer sequences are listed in Supplemental Table 3.

*FISH analysis.* FISH was performed using RNA Fluorescence In Situ Hybridization Kit (GenePharma) according to instructions. 18S and RP24-315D19.10 probes were designed and synthesized by GenePharma company and their 3’end was labeled with biotin. The sequences of 18S and RP24-315D19.10 probes are listed in Supplemental Table 5. In simple terms, the primary mESCs were inoculated on the sterile slide followed by artificially induced decidualization. mESCs and DSCs were then fixed with 4% paraformaldehyde for 15 minutes and permeated with 0.5% Triton X-100 for 15 minutes respectively. Next, cells were blocked with 1 x blocking buffer at 37 °C. Finally, 15 μl probe working solution (3 μl 1 μmol/l biotin probe + 1.5 μl 1 μmol/l SA-Cy3 + 10.5 μl PBS) and 85 μl hybrid buffer was mixed together and incubated with cells for 12-16 h at 37 °C. The next day, cells were washed with 0.1% Tween 20 and 2 x normal saline sodium citrate (SSC) buffer respectively. 18S and RP24-315D19.10 was measured under confocal microscopy. *RNA pull-down and mass spectrometry analysis.* To detect the interaction among RP24-315D19.10 and targeted proteins, we performed RNA-Protein pull-down assay by using Pierce™ Magnetic RNA-Protein Pull-Down Kit (Thermo Fisher Scientific). The biotin labeled RP24-315D19.10 and AS-RP24-315D19.10 probe was designed and synthesized by GenePharma company. The sequences of biotin labeled probes are listed in Supplemental Table 5. Briefly, the biotin labeled RP24-315D19.10 and AS-RP24-315D19.10 probes were bound to magnetic beads respectively before protein lysate form mESCs and DSCs was added. Beads were washed by adding the appropriate buffer, vortexed and separated on a magnetic stand. Finally, RNA binding proteins were separated by 10% SDS-PAGE gel, stained with Coomassie blue and then subjected to mass spectrometry analysis (Shanghai Applied Protein Technology Co., Ltd).

### RIP analysis

RNA immunoprecipitation was performed using the Magna RIP RNA-Binding Protein Immunoprecipitation Kit (Millipore, Billerica) following the manufacturer’s protocol. DSCs were lysed in complete RIP lysis buffer followed by centrifugation at 14,000 r.p.m for 10 min at 4 °C. Next, transfer appropriate amount of supernatant to RIP buffer containing beads, which have been conjugated with 5 μg of hnRNPA2B1 antibodies or control IgG, and the mixture was incubated with rotation at 4 °C for overnight. RNA was extracted from the immunoprecipitates and detected by RT-qPCR. The primer used for detecting RP24-315D19.10 is presented in Supplemental Table 3.

### Immunohistochemistry

Paraffin embedded tissue sections of human decidua were prepared on charged glass slides, followed by immunostained with hnRNPA2B1 antibodies (1:500; Abcam) according to directions of DAB color reagent kit (Zhongshan Biosciences Inc.).

### TUNEL staining

For cell apoptosis detection, we used Colorimetric TUNEL Apoptosis Assay Kit (Beyotime Biotechnology) according to the manufacturer’s protocol. Briefly, cells were fixed in 4% formaldehyde for 30 min at room temperature, washed twice in PBS and then treated with 0.3% Triton X-100 for 5 min. After double wash in PBS, samples were incubated with 0.3% H2O2 in PBS for 20 min at room temperature. The reaction mix was prepared according to manufacturer’s protocol. Staining was carried out at 37 °C for 60 min in dark. After staining termination, the cell apoptosis was observed and photographed under microscope.

### Human decidual sample collection

Human decidual tissues were collected when the participants were under surgery to terminate the pregnancy. Decidual tissues from normal women who underwent a legal termination of a healthy pregnancy (Ctrl) (n=6) and patients with spontaneous abortion (SA) (n=7) at gestational weeks 0–28 was obtained at the Chongqing Health Center for Women and Children. After the curettage all samples were immediately stored at −80 °C or formalin fixed for further analysis.

### Biolayer interferometry

The interaction between hnRNPA2B1 protein and lincRNA RP24-315D19.10 was verified by using the Octet QKe (ForteBio) essentially as described previously(63). Biotin-labeled RP24-315D19.10 was synthesized by GenePharma company, hnRNPA2B1 was purified by Biogot technology, co, Ltd (China). Streptavidin biosensors (ForteBio) were hydrated for 10 min prior to the experiment in sample diluents (PBS, 0.02% Tween 20). Biotin-labeled RP24-315D19.10 was at 50 nM, and purified hnRNPA2B1 was at 0 and 50 nM. The settings were as follows: initial base line for 120 s, loading for 600 s, base line for 60 s, association for 600 s, and dissociation for 600 s. *KD* values were generated by BLItz Pro software.

### Statistics

Statistical analysis was performed using SPSS 26.0 software, data were shown as mean ± standard deviation (SD) and presented with n denoting the sample size. The comparison between different groups was performed using Student’s *t* test. Pearson correlation analysis were also adopted in the course of analysis. *P* < 0.05 was considered to be statistically significant.

### Study approval

All our animal studies were approved by Ethics Committee of Chongqing Medical University. All included pregnant women signed informed consent, and the study was approved by the Ethics Committee of Chongqing Health Center for Women and Children (NO. yjkt20211213).

## Author contributions

JH and RG conceived, designed, and directed this research. LT performed most of the *in vivo* and *in vitro* experiments with the assistance of XC, YG, XY, CP, XM, FL and YZ. Clinical samples were collected and processed by YS. LT and RG performed the data analyses and finally prepared the manuscript. All authors read and approved the final manuscript. The order of the co-first authors was determined on the basis of their relative contributions.

## Acknowledgments

We are grateful for the financial support from the Special Professorship Project of Chongqing Medical University (2021-215).

## Notes

Conflict of interest: The authors have declared that no conflict of interest exists.

https://www.csbio.sjtu.edu.cn/bioinf/lncLocator/

https://www.ncbi.nlm.nih.gov/geo

http://service.tartaglialab.com/page/catrapid_group

### Competing Interest Statement

The authors have declared no competing interest.

