## Supplemental Figure 1-Supplemental Figure 7, Supplemental Table 1-Supplemental Table 5 for "lincRNA RP24-315D19.10 promotes endometrial decidualization via upregulation of hnRNPA2B1"

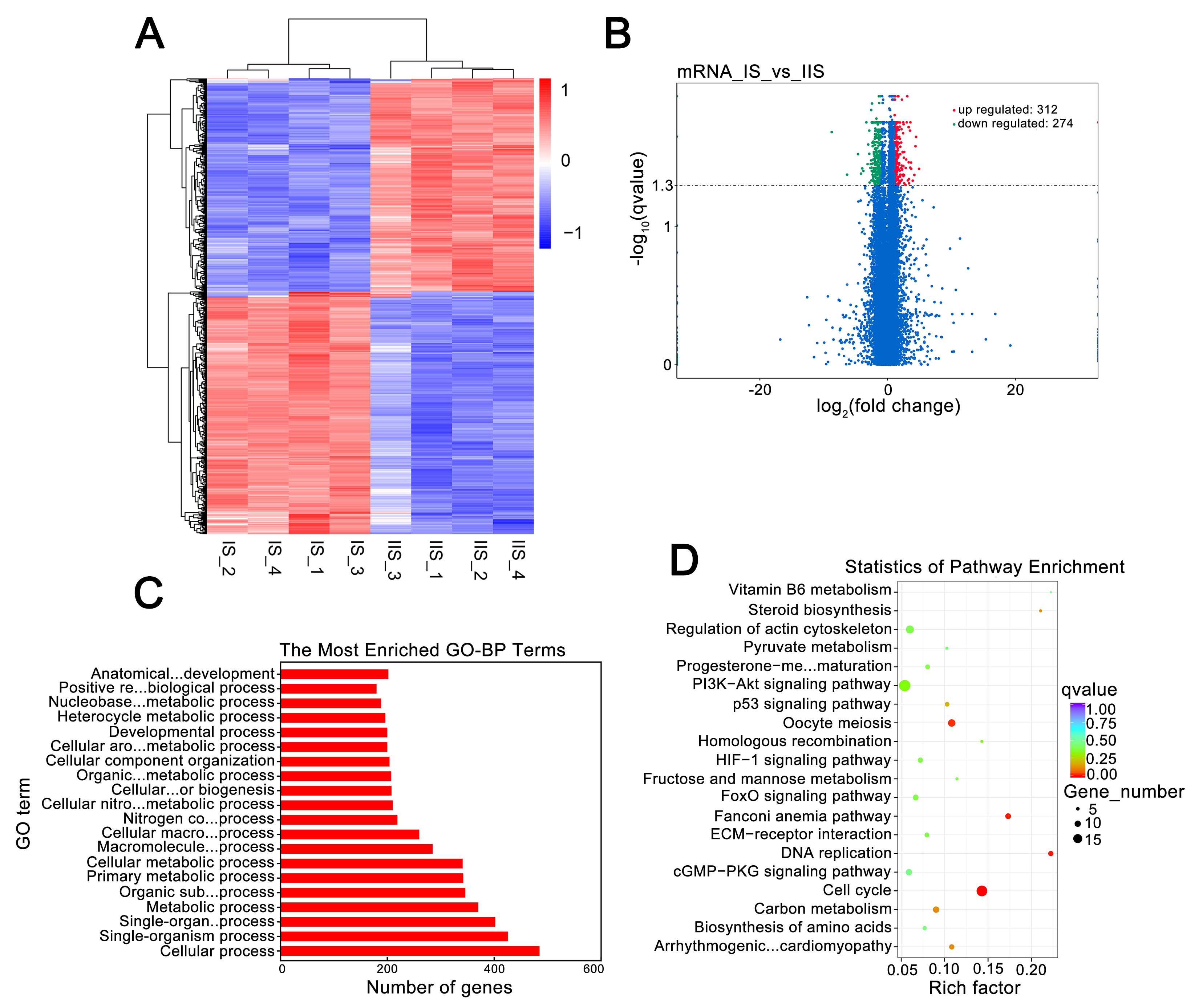

**Supplemental Figure** **1.**  **RNA-seq in mice endometrium at implantation and inter-implantation sites on the 6th day of pregnancy.** (**A**): The cluster heat map shows mRNAs with expression change fold ≥ 2 from RNA-Seq data (*P*<0.05).  Red color indicates high expression level, and blue color indicates low expression level. (**B**): Volcano plot shows the distribution of differentially expressed mRNAs. (**C**): Top 20 GO-BP terms enriched in differentially expressed mRNAs. (**D**): Top 20 KEGG pathways enriched in differentially expressed mRNAs. IS: implantation sites; IIS: inter-implantation sites.

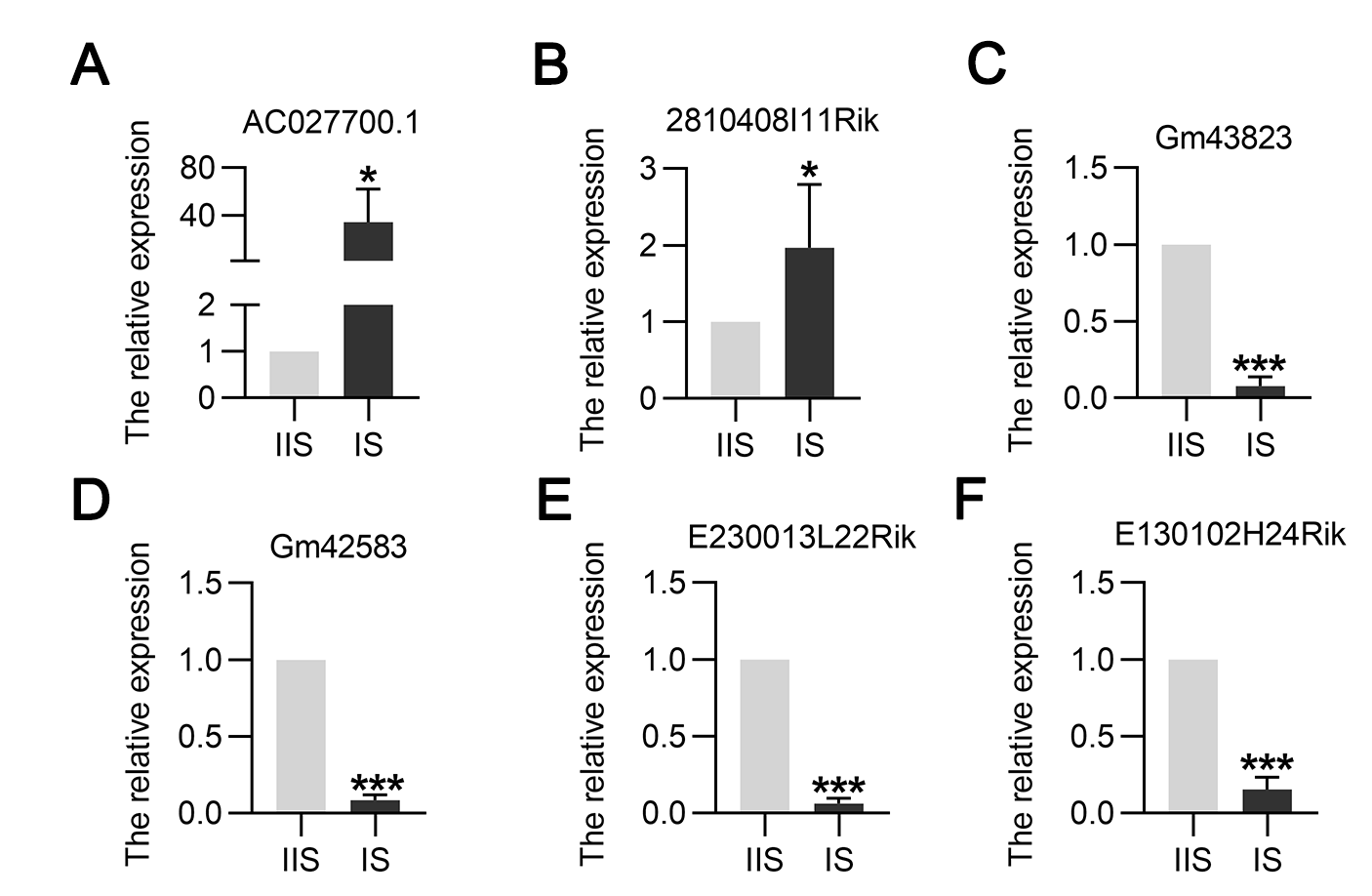

**Supplemental Figure 2. The expression levels of 6 dysregulated lncRNAs in mice endometrium on the 6th day of pregnancy.** (**A-F**): RT-qPCR detection of these 6 lncRNAs expression at the IIS and at the IS on 6th Day of pregnancy. IS: implantation sites; IIS: inter-implantation sites; **P* < 0.05, ****P* < 0.001.

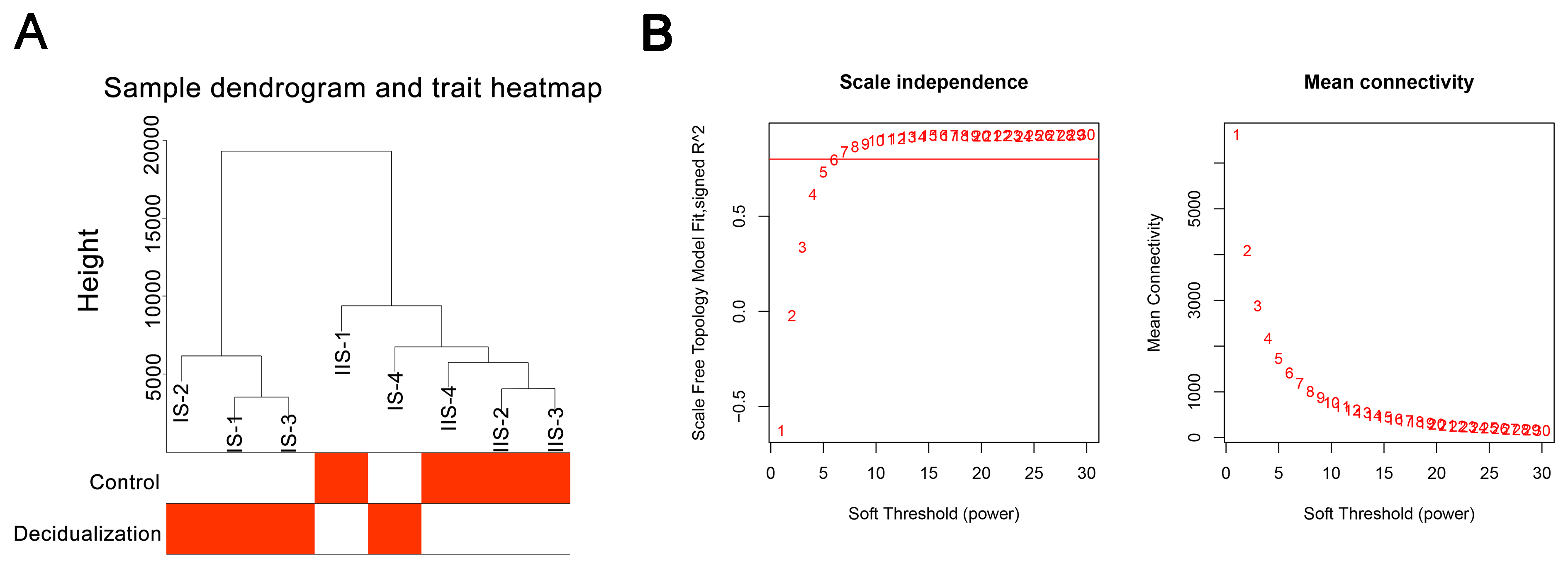

**Supplemental Figure 3. Identification of decidualization-associated hub lncRNAs by WGCNA.** (**A**): Hierarchical clustering dendrogram of samples and the corresponding trait heatmap. The red indicates a positive correlation between samples and decidualization, whereas white indicates a negative correlation. (**B**): Analysis of network topology for different soft‐threshold power (weighting coefficient, β). The left-hand image shows the impact of soft‐threshold power on the scale‐free topology fit index; the right-hand image displays average network connectivity under different soft‐threshold power. IS: implantation sites; IIS: inter-implantation sites.

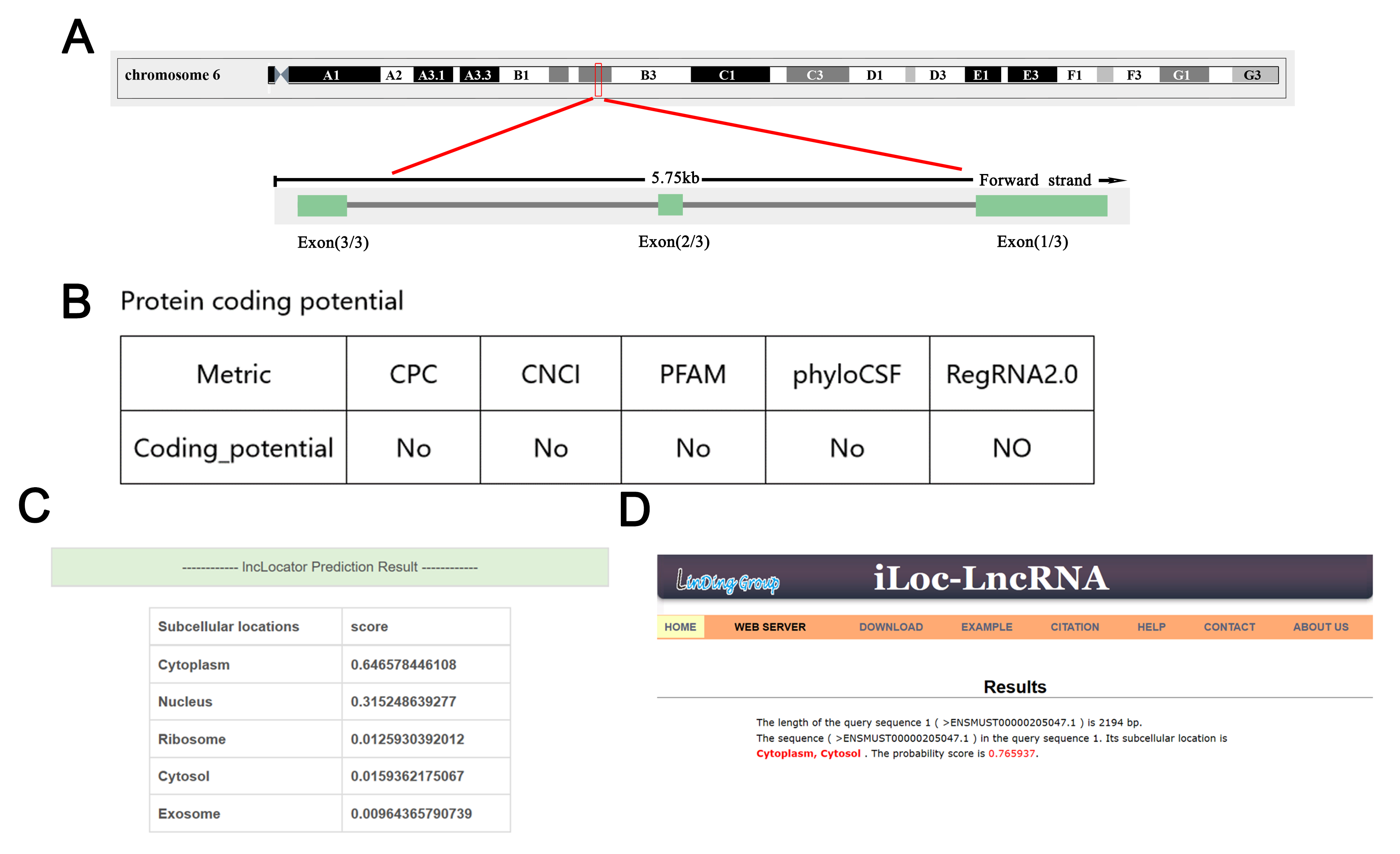

**Supplemental Figure 4. Predicted sub-cellular location of RP24-315D19.10.** (**A**): RP24-315D19.10 genome site; (**B**): the RP24-315D19.10 coding potential was predicted using 5 prediction tools. (**C-D**): The online software lncLocator and iLoc-LncRNA were used to predict the location of RP24-315D19.10.

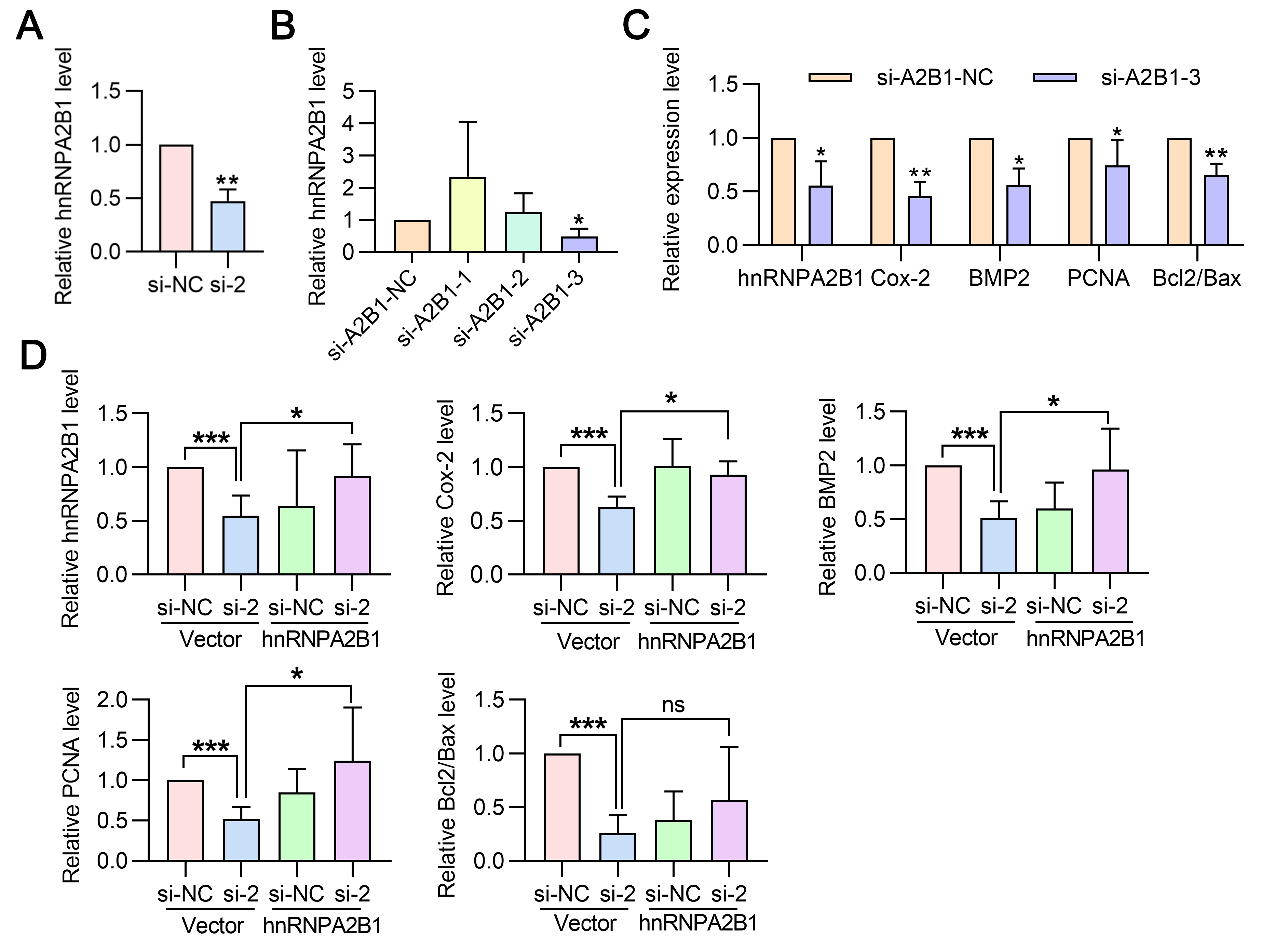

**Supplemental Figure 5. Statistical graphs for Figure 6.** (**A**): Western blot detection of hnRNPA2B1 protein expression level after RP24-315D19.10 silencing. Histogram showing the quantification of the results (n=4). (**B**): Western blot analysis indicates the hnRNPA2B1 expression after transfected with 3 different siRNAs targeting hnRNPA2B1 and negative control. Histogram showing the quantification of the results (n=3). (**C**): After hnRNPA2B1 silencing, the Cox-2, BMP2, PCNA and the ratio of Bcl2/Bax during decidualization were detected by western blot. Histogram showing the quantification of the results (n=4). (**D**): After overexpressed hnRNPA2B1, the Cox-2, BMP2, PCNA and the ratio of Bcl2/Bax in RP24-315D19.10 silencing mESCs during decidualization were detected by western blot. Histogram showing the quantification of the results(n=5). **P* < 0.05, ***P* < 0.01, ****P* < 0.001 compared with controls; data are presented as mean ± SD. BMP2: bone morphogenic protein 2; Cox-2: Cyclooxygenase-2; PCNA: proliferating cell nuclear antigen; Bcl2: B-cell lymphoma 2.

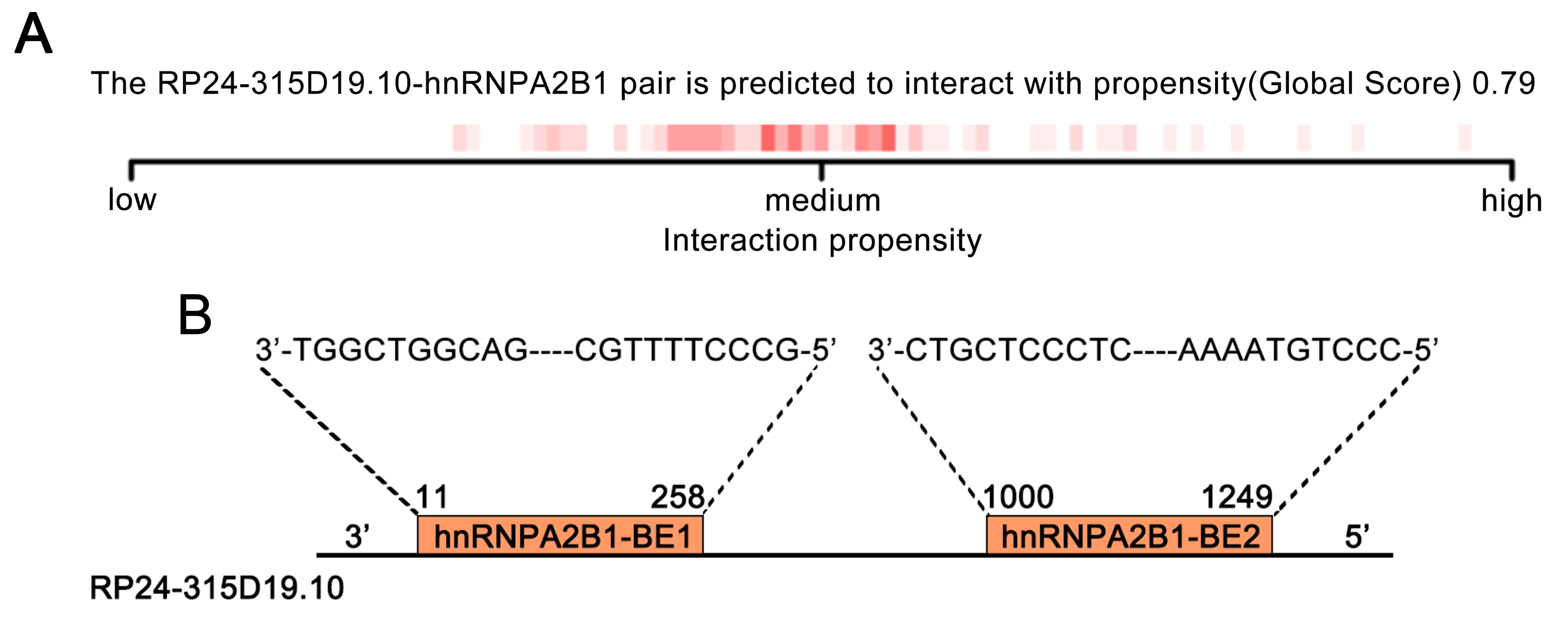

**Supplemental Figure 6. Recognition of RP24-315D19.10/hnRNPA2B1 binding regions.** (**A**): catRAPID interactions with large RNAs module prediction of the interaction propensity between hnRNPA2B1 protein and RP24-315D19.10. (**B**): Alignment of the hnRNPA2B1-binding element sequences of the RP24-315D19.10 sequence.

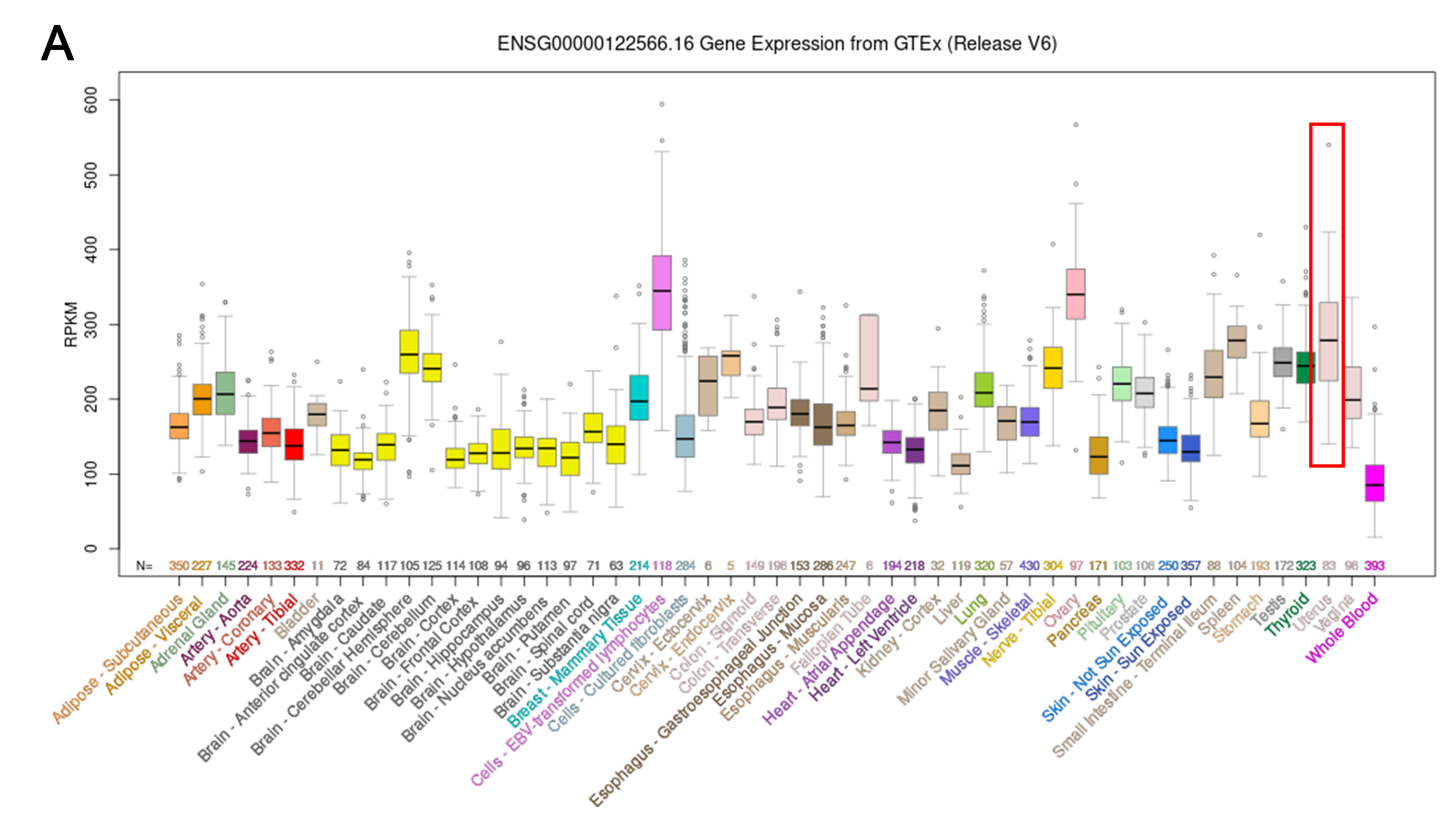

**Supplemental Figure 7. hnRNPA2B1 expression in human uterine tissues.** (**A**): Median human hnRNPA2B1 expression levels in 52 tissues and 2 cell lines, based on RNA-seq data from the GTEx final data release (V8, August 2019).

**Supplemental Table.1 The 155 hub RNAs in blue module**

| RNAs | GS.Decidulization | *P*_value | MMblue | *P*_value |
| --- | --- | --- | --- | --- |
| Gm4258 | 0.985411 | 7.68E-06 | 0.99439 | 4.40E-07 |
| RP24-315D19.10 | 0.984081 | 9.96E-06 | 0.99094 | 1.85E-06 |
| Gm26737 | 0.98483 | 8.63E-06 | 0.9859 | 6.93E-06 |
| 2010204K13Rik | 0.99596 | 1.64E-07 | 0.980468 | 1.84E-05 |
| Bzw1 | 0.987884 | 4.41E-06 | 0.997465 | 4.06E-08 |
| Hjurp | 0.984778 | 8.72E-06 | 0.980013 | 1.97E-05 |
| Cacybp | 0.980898 | 1.72E-05 | 0.988019 | 4.26E-06 |
| Parp1 | 0.99181 | 1.36E-06 | 0.98117 | 1.65E-05 |
| Cenpf | 0.989595 | 2.79E-06 | 0.989557 | 2.82E-06 |
| Nek2 | 0.980786 | 1.75E-05 | 0.996203 | 1.36E-07 |
| Nedd1 | 0.98263 | 1.29E-05 | 0.985352 | 7.77E-06 |
| Cct2 | 0.992547 | 1.03E-06 | 0.981258 | 1.62E-05 |
| Cdk4 | 0.990437 | 2.17E-06 | 0.993042 | 8.38E-07 |
| Prim1 | 0.992752 | 9.47E-07 | 0.98493 | 8.46E-06 |
| Cs | 0.988273 | 4.00E-06 | 0.992139 | 1.21E-06 |
| Fignl1.1 | 0.984195 | 9.75E-06 | 0.990649 | 2.03E-06 |
| Spdl1 | 0.992007 | 1.27E-06 | 0.983674 | 1.07E-05 |
| Hmmr | 0.982685 | 1.28E-05 | 0.994069 | 5.19E-07 |
| Hnrnpab.2 | 0.985372 | 7.74E-06 | 0.98287 | 1.24E-05 |
| Vdac1 | 0.98183 | 1.48E-05 | 0.997211 | 5.41E-08 |
| Eif4a1 | 0.985383 | 7.72E-06 | 0.984496 | 9.21E-06 |
| Rpa1 | 0.982358 | 1.35E-05 | 0.993735 | 6.12E-07 |
| Appbp2 | 0.985756 | 7.15E-06 | 0.987336 | 5.03E-06 |
| Prr11 | 0.987576 | 4.75E-06 | 0.996486 | 1.08E-07 |
| Acly.1 | 0.987202 | 5.19E-06 | 0.986614 | 5.94E-06 |
| Nsf | 0.984065 | 1.00E-05 | 0.985411 | 7.68E-06 |
| Kpna2 | 0.991935 | 1.30E-06 | 0.990217 | 2.32E-06 |
| Ube2o | 0.980079 | 1.95E-05 | 0.985793 | 7.09E-06 |
| Tk1 | 0.983499 | 1.11E-05 | 0.988345 | 3.92E-06 |
| Birc5 | 0.995132 | 2.87E-07 | 0.997038 | 6.48E-08 |
| Pole2 | 0.981581 | 1.54E-05 | 0.987045 | 5.38E-06 |
| Sav1 | 0.986071 | 6.69E-06 | 0.988959 | 3.34E-06 |
| Psma3 | 0.985257 | 7.92E-06 | 0.989393 | 2.96E-06 |
| Ttc7b | 0.984037 | 1.00E-05 | 0.98909 | 3.22E-06 |
| Hist1h2ap | 0.987214 | 5.18E-06 | 0.990675 | 2.01E-06 |
| Hist1h3c | 0.982061 | 1.42E-05 | 0.993703 | 6.21E-07 |
| Hist1h1a | 0.982351 | 1.36E-05 | 0.993675 | 6.30E-07 |
| Cenpp | 0.986928 | 5.53E-06 | 0.989194 | 3.13E-06 |
| Pdlim7 | 0.988248 | 4.02E-06 | 0.986066 | 6.69E-06 |
| Ak6.1 | 0.9883 | 3.97E-06 | 0.9892 | 3.12E-06 |
| Vcl | 0.98649 | 6.10E-06 | 0.984535 | 9.14E-06 |
| Nt5dc2 | 0.990631 | 2.04E-06 | 0.99523 | 2.70E-07 |
| Mapk1ip1l | 0.984701 | 8.85E-06 | 0.986127 | 6.61E-06 |
| Dlgap5 | 0.989291 | 3.05E-06 | 0.997235 | 5.27E-08 |
| Pbk | 0.986159 | 6.56E-06 | 0.995866 | 1.76E-07 |
| Cdca2 | 0.993653 | 6.36E-07 | 0.98395 | 1.02E-05 |
| Cdca2.1 | 0.992893 | 8.93E-07 | 0.988053 | 4.22E-06 |
| Rad21 | 0.991408 | 1.58E-06 | 0.995735 | 1.93E-07 |
| Sh3bp1 | 0.984141 | 9.85E-06 | 0.985473 | 7.58E-06 |
| Ttll12 | 0.984234 | 9.68E-06 | 0.984531 | 9.15E-06 |
| Cbx5 | 0.98904 | 3.26E-06 | 0.985974 | 6.83E-06 |
| Trap1 | 0.983365 | 1.14E-05 | 0.990948 | 1.84E-06 |
| Mcm4 | 0.991162 | 1.71E-06 | 0.989111 | 3.20E-06 |
| Ccdc58 | 0.99002 | 2.47E-06 | 0.995665 | 2.03E-07 |
| Phldb2 | 0.988793 | 3.49E-06 | 0.987538 | 4.79E-06 |
| C330027C09Rik | 0.991662 | 1.44E-06 | 0.985714 | 7.21E-06 |
| Mis18a | 0.99161 | 1.47E-06 | 0.986711 | 5.81E-06 |
| Wrb | 0.990123 | 2.39E-06 | 0.988213 | 4.06E-06 |
| Mrpl18 | 0.987942 | 4.34E-06 | 0.995084 | 2.96E-07 |
| Srsf3 | 0.994506 | 4.13E-07 | 0.990689 | 2.00E-06 |
| Nrm | 0.993534 | 6.73E-07 | 0.996322 | 1.24E-07 |
| Sgo1 | 0.987997 | 4.28E-06 | 0.991934 | 1.30E-06 |
| Emilin2 | 0.980108 | 1.94E-05 | 0.988679 | 3.60E-06 |
| Gm10093 | 0.981015 | 1.69E-05 | 0.983991 | 1.01E-05 |
| Hnrnpll | 0.990251 | 2.30E-06 | 0.991343 | 1.61E-06 |
| Pkdcc | 0.986167 | 6.55E-06 | 0.989243 | 3.09E-06 |
| Hspa9 | 0.98485 | 8.60E-06 | 0.987266 | 5.11E-06 |
| Eno1b | 0.988351 | 3.92E-06 | 0.992806 | 9.26E-07 |
| Lmnb1 | 0.990437 | 2.17E-06 | 0.982655 | 1.29E-05 |
| Ppp1r14b | 0.984523 | 9.16E-06 | 0.980232 | 1.90E-05 |
| Kif11 | 0.987343 | 5.02E-06 | 0.988653 | 3.62E-06 |
| Entpd7 | 0.984173 | 9.79E-06 | 0.981739 | 1.50E-05 |
| Prrx2 | 0.983842 | 1.04E-05 | 0.996826 | 7.97E-08 |
| Fam129b | 0.980774 | 1.75E-05 | 0.991506 | 1.52E-06 |
| Mtx2 | 0.989354 | 2.99E-06 | 0.991345 | 1.61E-06 |
| Api5 | 0.989151 | 3.17E-06 | 0.980435 | 1.84E-05 |
| Pdhx | 0.986969 | 5.48E-06 | 0.990278 | 2.28E-06 |
| Kif18a | 0.994176 | 4.92E-07 | 0.982358 | 1.35E-05 |
| Bub1b | 0.996226 | 1.34E-07 | 0.994752 | 3.60E-07 |
| Rad51 | 0.985092 | 8.19E-06 | 0.986156 | 6.56E-06 |
| Ncaph | 0.989887 | 2.57E-06 | 0.993752 | 6.07E-07 |
| Bub1 | 0.991678 | 1.43E-06 | 0.987243 | 5.14E-06 |
| Ckap2l | 0.991057 | 1.78E-06 | 0.987841 | 4.45E-06 |
| Gins1 | 0.984007 | 1.01E-05 | 0.985407 | 7.68E-06 |
| Fkbp1a | 0.9908 | 1.93E-06 | 0.989074 | 3.23E-06 |
| Ctnnbl1 | 0.9879 | 4.39E-06 | 0.99402 | 5.32E-07 |
| Ywhab | 0.99022 | 2.32E-06 | 0.991072 | 1.77E-06 |
| Cebpb | 0.981576 | 1.54E-05 | 0.983569 | 1.10E-05 |
| Prkci | 0.991714 | 1.41E-06 | 0.987803 | 4.49E-06 |
| Ccna2 | 0.988217 | 4.05E-06 | 0.986705 | 5.82E-06 |
| Smc4 | 0.997393 | 4.42E-08 | 0.99122 | 1.68E-06 |
| Pmf1 | 0.987248 | 5.13E-06 | 0.984753 | 8.76E-06 |
| Efna4 | 0.980315 | 1.88E-05 | 0.983467 | 1.12E-05 |
| Cenpe | 0.998161 | 1.55E-08 | 0.988744 | 3.53E-06 |
| H2afz | 0.980348 | 1.87E-05 | 0.994054 | 5.23E-07 |
| Depdc1a | 0.980433 | 1.85E-05 | 0.981943 | 1.45E-05 |
| Frmpd1 | 0.988952 | 3.34E-06 | 0.991959 | 1.29E-06 |
| Anp32b | 0.987966 | 4.32E-06 | 0.995349 | 2.51E-07 |
| Ror1 | 0.983185 | 1.17E-05 | 0.992789 | 9.33E-07 |
| Kif2c | 0.992344 | 1.12E-06 | 0.995893 | 1.73E-07 |
| Rpa2 | 0.986538 | 6.04E-06 | 0.990498 | 2.13E-06 |
| Nudc | 0.980598 | 1.80E-05 | 0.986182 | 6.53E-06 |
| Mrto4 | 0.980977 | 1.70E-05 | 0.988044 | 4.23E-06 |
| Vamp3 | 0.990573 | 2.08E-06 | 0.986184 | 6.53E-06 |
| Ncapg | 0.9811 | 1.66E-05 | 0.985659 | 7.30E-06 |
| Ugdh | 0.99223 | 1.17E-06 | 0.990168 | 2.36E-06 |
| Nipal1 | 0.981255 | 1.62E-05 | 0.990817 | 1.92E-06 |
| Paqr3 | 0.984358 | 9.46E-06 | 0.98913 | 3.18E-06 |
| Kmt5a | 0.982541 | 1.31E-05 | 0.995364 | 2.48E-07 |
| Ran | 0.984505 | 9.19E-06 | 0.990059 | 2.44E-06 |
| Limk1 | 0.990937 | 1.85E-06 | 0.983242 | 1.16E-05 |
| Aoc1.1 | 0.982957 | 1.22E-05 | 0.993885 | 5.69E-07 |
| Fkbp9 | 0.984666 | 8.91E-06 | 0.981833 | 1.48E-05 |
| Ncapd2 | 0.990082 | 2.42E-06 | 0.988707 | 3.57E-06 |
| Rdh13 | 0.982693 | 1.28E-05 | 0.988258 | 4.01E-06 |
| U2af2.1 | 0.984869 | 8.56E-06 | 0.989797 | 2.63E-06 |
| Sae1 | 0.992109 | 1.22E-06 | 0.993751 | 6.07E-07 |
| E2f8.1 | 0.983714 | 1.07E-05 | 0.98499 | 8.36E-06 |
| Fanci | 0.99381 | 5.90E-07 | 0.984053 | 1.00E-05 |
| Ddias | 0.985355 | 7.77E-06 | 0.980863 | 1.73E-05 |
| Rras2 | 0.98454 | 9.13E-06 | 0.994529 | 4.08E-07 |
| Psma1 | 0.981454 | 1.57E-05 | 0.989386 | 2.97E-06 |
| Eif3c | 0.983566 | 1.10E-05 | 0.981793 | 1.49E-05 |
| Hirip3 | 0.985126 | 8.13E-06 | 0.986577 | 5.99E-06 |
| Kif22 | 0.986567 | 6.00E-06 | 0.998313 | 1.20E-08 |
| Shcbp1 | 0.995763 | 1.89E-07 | 0.996406 | 1.16E-07 |
| Ckap2 | 0.994636 | 3.84E-07 | 0.982139 | 1.41E-05 |
| Asf1b | 0.990916 | 1.86E-06 | 0.99526 | 2.65E-07 |
| Polr2c | 0.993408 | 7.13E-07 | 0.989174 | 3.15E-06 |
| Plekhg4.2 | 0.980636 | 1.79E-05 | 0.988363 | 3.91E-06 |
| Rfwd3 | 0.984866 | 8.57E-06 | 0.991199 | 1.69E-06 |
| Cfdp1 | 0.988944 | 3.35E-06 | 0.984712 | 8.83E-06 |
| Cdt1 | 0.980054 | 1.95E-05 | 0.995931 | 1.68E-07 |
| Dnmt1 | 0.990246 | 2.30E-06 | 0.981855 | 1.47E-05 |
| Tmed1 | 0.986945 | 5.51E-06 | 0.989581 | 2.81E-06 |
| Anln | 0.991764 | 1.39E-06 | 0.99197 | 1.29E-06 |
| H2afx | 0.984631 | 8.97E-06 | 0.996323 | 1.24E-07 |
| Acat1 | 0.982718 | 1.27E-05 | 0.993105 | 8.15E-07 |
| Idh3a | 0.98533 | 7.81E-06 | 0.996283 | 1.28E-07 |
| Pkm | 0.986874 | 5.60E-06 | 0.990587 | 2.07E-06 |
| Smad6 | 0.993381 | 7.21E-07 | 0.992338 | 1.12E-06 |
| Pclaf | 0.992216 | 1.17E-06 | 0.997955 | 2.14E-08 |
| Ccnb2 | 0.986033 | 6.74E-06 | 0.995027 | 3.06E-07 |
| Ttk | 0.990889 | 1.88E-06 | 0.989185 | 3.14E-06 |
| Pfkfb4 | 0.981397 | 1.59E-05 | 0.987464 | 4.88E-06 |
| Ctdspl | 0.991567 | 1.49E-06 | 0.985737 | 7.18E-06 |
| Clcn5 | 0.993517 | 6.78E-07 | 0.987514 | 4.82E-06 |
| Suv39h1 | 0.987813 | 4.48E-06 | 0.996779 | 8.34E-08 |
| Efnb1 | 0.9814 | 1.59E-05 | 0.988019 | 4.26E-06 |
| Nono | 0.988828 | 3.46E-06 | 0.981632 | 1.53E-05 |
| Pgk1 | 0.988451 | 3.82E-06 | 0.993405 | 7.14E-07 |
| Mcm3 | 0.991957 | 1.29E-06 | 0.99384 | 5.82E-07 |
| Prim2 | 0.9827 | 1.28E-05 | 0.988261 | 4.01E-06 |
| Hspd1 | 0.987615 | 4.71E-06 | 0.991822 | 1.36E-06 |
| Hspe1 | 0.982593 | 1.30E-05 | 0.990524 | 2.11E-06 |

**Supplemental Table.2 The top 20 predicted binding regions of RP24-315D19.10**

| # | Protein | RNA region | Interaction Propensity |
| --- | --- | --- | --- |
| 1 | HnRNPA2B1 | ENSMUST00000205047.1-45-241 | 34.9 |
| 2 | HnRNPA2B1 | ENSMUST00000205047.1-66-258 | 33.88 |
| 3 | HnRNPA2B1 | ENSMUST00000205047.1-11-198 | 29.52 |
| 4 | HnRNPA2B1 | ENSMUST00000205047.1-69-242 | 28.92 |
| 5 | HnRNPA2B1 | ENSMUST00000205047.1-125-258 | 25.41 |
| 6 | HnRNPA2B1 | ENSMUST00000205047.1-15-177 | 25.33 |
| 7 | HnRNPA2B1 | ENSMUST00000205047.1-113-218 | 24.5 |
| 8 | HnRNPA2B1 | ENSMUST00000205047.1-977-1171 | 22.79 |
| 9 | HnRNPA2B1 | ENSMUST00000205047.1-986-1188 | 22.27 |
| 10 | HnRNPA2B1 | ENSMUST00000205047.1-81-192 | 19.89 |
| 11 | HnRNPA2B1 | ENSMUST00000205047.1-79-191 | 18.84 |
| 12 | HnRNPA2B1 | ENSMUST00000205047.1-1065-1191 | 17.31 |
| 13 | HnRNPA2B1 | ENSMUST00000205047.1-1057-1169 | 15.17 |
| 14 | HnRNPA2B1 | ENSMUST00000205047.1-1000-1171 | 14.3 |
| 15 | HnRNPA2B1 | ENSMUST00000205047.1-1096-1238 | 13.69 |
| 16 | HnRNPA2B1 | ENSMUST00000205047.1-1036-1171 | 12.83 |
| 17 | HnRNPA2B1 | ENSMUST00000205047.1-1039-1169 | 12.79 |
| 18 | HnRNPA2B1 | ENSMUST00000205047.1-1063-1243 | 12.24 |
| 19 | HnRNPA2B1 | ENSMUST00000205047.1-1077-1249 | 10.49 |
| 20 | HnRNPA2B1 | ENSMUST00000205047.1-303-479 | 8.84 |

**Supplemental Table.3 The sequences of specific primers**

| Target | Primer sequence（5’ to 3’) |
| --- | --- |
| Gm4258 | F: GCCTGAGCACAAGTAGCGAGTTC |
|  | R: CGGTGCAGGTGCAGAGTCTTG |
| RP24-315D19.10 | F: TTGCTTGCTGTGGCTGCTTCTC |
|  | R: GGCAGAACACTCCGACACATACAG |
| 2010204K13Rik | F: GCGGTGCAGTTTCAGATGAC |
|  | R: AACAGCTCTTTGCAGGCTTTC |
| Gm26737 | F: CGTCTAACCAAGGAAGAAGCAAG |
|  | R: CGCTGTCACTTTAGAGATGTTTCAC |
| β-actin | F: GTGCTATGTTGCTCTAGACTTCG |
|  | R: ATGCCACAGGATTCCATACC |
| Dtprp | F: TACCCACGTAAGGTCATC |
|  | R: CTCAGAGCCAGAAATCA |
| U1 | F: TTTCCCAGGGCGAGGCTCAC |
|  | R: TGCAGTCGAGTTTCCCGCATTTG |
| U6 | F: GCTCGCTTCGGCAGCACATATAC |
|  | R: CGAATTTGCGTGTCATCCTTGCG |
| GAPDH | F: CAGGAGGCATTGCTGATGAT |
|  | R: GAAGGCTGGGGCTCATTT |
| Linc00276 | F: GGCCAAGAAACCAACCAGTG |
|  | R: TCACCACTGGCACTTCCATTT |
| TRAM2-AS1 | F: GACCTCCTGCGAACAACCATCAC |
|  | R: AAGGCTTAACTGGGGCTGGAAAATC |
| AC027700.1 | F: CGGAAGTTCAGCAGAAGGAAGAGG |
|  | R: GCCCACACCTTATTCACCAAGACC |
| 2810408I11Rik | F: AGTGTGGTCCAGAGAGCAGAGC |
|  | R: ACAGCATGTGATGAGAGCCAGTTG |
| Gm43823 | F: ACAGGGCAGAACCTCTTAGTAGC |
|  | R: GGGAGTAAAAGAAAGTGCGGT |
| Gm42583 | F: CAACAAGAATTGCTGAAGGGCT |
|  | R: ACGGTGATTATGAGAAGCACGAC |
| E230013L22Rik | F: CCAACGCTCTCTGACTCAGT |
|  | R: AGTACGACTCTCGCTTCTTAGGTG |
| E130102H24Rik | F: ACACACTTCACACCGCTCCTG |
|  | R: GCTGCTACATTGTTTTGCTCCTT |

**Supplemental Table.4 The sequences of specific siRNAs**

| Gene | NO. | siRNA（5’ to 3’) |
| --- | --- | --- |
| RP24-315D19.10 | Si-1 | CUCCUGUUGAUGGCAUUAUTT |
|  | Si-2 | CCUCAUGGGUUGUGAAUAATT |
|  | Si-3 | GGGUUUCACUCUGCCUUAUTT |
|  | Si-NC | UUCUCCGAACGUGUCACGUTT |
| HnRNPA2B1 | si-hnRNPA2B1-1 | GGGCUUCACUGUAUAAAUATT |
|  | si-hnRNPA2B1-2 | GGACCAGGAAGCAACUUUATT |
|  | si-hnRNPA2B1-3 | GCCAGGAUCAUGGUGUAAUTT |
|  | si-hnRNPA2B1-NC | UUCUCCGAACGUGUCACGUTT |

**Supplemental Table.5 The sequences of specific probes**

| Target | Application | Probe sequence（5’ to 3’) |
| --- | --- | --- |
| RP24-315D19.10 | FISH | TGA+TGATGGC+TGTAAAGCA+TAGGGCAA |
| 18S |  | CTGCCTTCCTTGGATGTGGTAGCCGTTTC |
| RP24-315D19.10 | RNA-Pulldown | UGAUGAUGGCUGUAAAGCAUAGGGCAA |
| Antisense |  | UUGCCCUAUGCUUUACAGCCAUCAUCA |

**The sequence of RP24-315D19.10:**

>ENSMUST00000205047.1

ACAGAATGAGTGGCTGGCAGATTGGTTATGAAAGCATGGCCCCCGACGGAGAGGGTCGAGATCCCAGGACACCTAGACTCAGCAGAGTCTCTACAGGCACCCCGCCTCTCTCCAGCGCCCCAGCTGGAGAAGGGGCCGGCACCCCCACCCCCCACCCCCGCCCACCCGCGATGCCTAAAGAGGAAGCTGCGTGGCCCCTTGCTTGCTGTGGCTGCTTCTCCCAGGTACAGGAGGCACGTGCGCTTCTGCGTTTTCCCGACAGGCGGCGCTATTCTAATGCATTTACTTACTGTATGTGTCGGAGTGTTCTGCCTGCATAAATGTCACATATACATGTGTGGCTGGAGCCCATGGAGGTCAGAAGAGAGAAGGATAACCCAGGCTGCTCTCTACGTCGCAAGAGCTTGAGGACCAGCCCAGCTGCAAAGCATCTGTCATGTCATGTTCTGTGAACTCAGATGCCATCCTGAGAGTCACCATTCATCCCAGCTCCAGCATCATTTAATGAACTTTTACAAATGTTTCCATTGGGTACCCCAAGTTCCCCTGAGGATGCAGAGTGCTTCATCACCTGTGGGGAGCACTCTCAGACCCCTTGGCAGTAAGTCACTTCTCCTACTCTACCCACCAGAGACAAACACTTGTGTCTTCACTTCCTGCTGAAAATGAATTAATAGCCCTTCAACTTTTGGCTCTGGCTTCTCCTACTTAGCAAAGCATCACTGGGATTTGTCCACACTATGTCAGTATCATCTTTTTTAAATTGCTGAGTAGTATTCAAGCTAGACAAGCCTTAGTCAACTTATCCTGCCCCCTCCTGTTGATGGCATTATGATCTTTGTTGCTGACATTGTTTATCCCTTGGCTTGTAATATCCTCCTTGTCATTGACCTGTGGACCCCACACAAAATCACACCCATAAGCCTTCCAAGACGCAGTGCGAAGGCTGGGGCTTTGAAAAACACTTTGTAATCCACTGCCACACCTCAGCTCAGGGGACTGCTCCCTCCTCCCTACTTGTTCATACTTTGACCTTTGTCCTTGACTGTTTTATGATCCTCACTTGTTTATAGATTTGACTCCTATTACTGCAGCCCACACCGCAGAGGGCAGGCACCAGGGGGTTTCACTCTGCCTTATACCTGGAGCTGCCTGGAGCAGGAGGGGCACAGGCTTGGCAAGCTGAAACCAATCCTACCTTTTCCACCATTAACATAATGATCCTCATGGGTTGTGAATAAAATGTCCCCACAAAGCTCTTGGCATGGAGCAAGTTTCTGAAACAGGGGGATAAAACTTACTTGTGGAAGACACAGAGGAGCCTGTGATAAAGTTAGTCACATGCCATGAAGCCTGGGTTTCTTTTCATTCTTATAGTACAGAAAGATTTGCTTTGTTTCAAAAACAGACATGACCTGGAGAGCCAGATAAGCAGGGAGTTCCACTTCCCATTTACCTTACTTTGCTTTTTATTGCTGTGATAAACACCAGGGCCCAAAGCAAGTAGGGCCTGAGAGGGTTTGCTTTGCCCTATGCTTTACAGCCATCATCAGAAGACAAGACAGGAACTCAAGTGGGGCAGGAACCTGGGGAGTTGAAGAAGAGGCCATGGAGGAATTCTGCTTACTCGCTTGCTTTCTGTGGTACATTCAACTTGACCTTTTATACAACCCATGATCACCTGCCCAGGAGTGGGTCCACCCCTATCAATTATTGATTAGAAAATTCCCCACAAGCTCTCCTAAGGGCCAGTGAAAAGACTAGTCAACTGAGGGTCTCTCTTCCCAGATGACTGTAGTGTGTGTCAAGCTGACACAATGTTCCAGGCTAACTGGCACAACTGCTCTCTTGACAGATTGACACAGGGACACATCATTGTTAAACTGCAACTTTTACTTTCTTGGATATTACCAGTATTTCATGGTAACACCTCGGTATAAAACATATCATAATCTTAAAAGTCCCATAGTCTTTTACATTTTAAAAGTCCATGTTCCTGCCTGGCAAGATGACTCAGGGGTTAAGAGCACTGACTGCTCTTCCAGAGGTTCTGAGTTCGATTCCCAGCAACCACATAGTGGTTTACAACCATCTGTAATGGGATCCAATGCCCTCTTCTGGTGTGTCTGAAGACAGCAACGGTATACTCACATATGTAAAATAAATAAATAAATCTTTAAATTCCTGCTCCCCT
